# Onecut factors and Pou2f2 regulate the distribution of V2 interneurons in the mouse developing spinal cord

**DOI:** 10.1101/413054

**Authors:** Audrey Harris, Gauhar Masgutova, Amandine Collin, Mathilde Toch, Maria Hidalgo-Figueroa, Benvenuto Jacob, Lynn M. Corcoran, Cédric Francius, Frédéric Clotman

## Abstract

Acquisition of proper neuronal identity and position is critical for the formation of neural circuits. In the embryonic spinal cord, cardinal populations of interneurons diversify into specialized subsets and migrate to defined locations within the spinal parenchyma. However, the factors that control interneuron diversification and migration remain poorly characterized. Here, we show that the Onecut transcription factors are necessary for proper diversification and distribution of the V2 interneurons in the developing spinal cord. Furthermore, we uncover that these proteins restrict and moderate the expression of spinal isoforms of *Pou2f2*, a transcription factor known to regulate B-cell differentiation. By gain- or loss-of-function experiments, we show that Pou2f2 contribute to regulate the position of V2 populations in the developing spinal cord. Thus, we uncovered a genetic pathway that regulates the diversification and the distribution of V2 interneurons during embryonic development.

**Significance statement:** In this study, we identify the Onecut and Pou2f2 transcription factors as regulators of spinal interneuron diversification and migration, two events that are critical for proper CNS development.

## Introduction

Neuronal migration is a critical feature of CNS development. It enables neurons to reach an adequate location in the nervous parenchyma and to properly integrate into neural circuits. The mechanisms that regulate neuronal migration have been extensively studied in the developing brain, particularly for cortical interneurons (INs) (Barber and Pierani, 2016, Guo and Anton, 2014). In contrast, the factors that control IN migration in the developing spinal cord remain almost totally unknown.

In the embryonic spinal cord, distinct neuronal populations are generated from different progenitor domains orderly distributed along the dorso-ventral axis of the ventricular zone. These progenitors produce motor neurons and multiple populations of ventral or dorsal INs (Lai et al., 2016, Lu et al., 2015). Although spinal IN populations do not organize into columns along the anteroposterior axis of the spinal cord, they each migrate according to a stereotyped pattern and settle down at specific focused or diffuse locations in the spinal parenchyma (Grossmann et al., 2010). Recent studies demonstrated that proper neuronal distribution is critical for adequate formation of spinal circuits. Indeed, the clustering and dorso-ventral settling position of motor neuron pools critically pattern sensory input specificity (Surmeli et al., 2011). Position of dorsal INs along the medio-lateral axis in lamina V determines their connectivity with sensory afferents (Hilde et al., 2016) while extensor and flexor premotor INs segregate along the medio-lateral axis of the spinal cord (Tripodi et al., 2011). Positional distinctions among premotor INs additionally correlate with their output to different motor columns (Goetz et al., 2015) and differential distribution of V1 IN subsets constrain patterns of input from sensory and from motor neurons (Bikoff et al., 2016). Consistently, distinct ventral IN subsets are differentially distributed along the anteroposterior axis of the spinal cord (Bikoff et al., 2016, Francius et al., 2013, Hayashi et al., 2018) and integrate into specific local microcircuit modules (Hayashi et al., 2018). However, the molecular mechanisms that regulate proper distribution of spinal INs remain elusive.

During their migration, cardinal populations of spinal neurons undergo progressive diversification into distinct subsets that exert specific functions in spinal circuits (Catela et al., 2015, Lai et al., 2016, Lu et al., 2015). For example, V2 INs divide into major V2a and V2b and minor V2c and V2d populations characterized by the expression of Chx10, Gata3, Sox1 and Shox2, respectively. V2a and V2d are excitatory neurons that participate in left-right alternation at high speed and contribute to rhythmic activation of locomotor circuits, respectively (Crone et al., 2008, Dougherty and Kiehn, 2010, Dougherty et al., 2013). V2b cells are inhibitory INs that participate in alternation of flexor vs extensor muscle contraction (Britz et al., 2015). As observed for V1 INs (Bikoff et al., 2016), V2 cells further diversify into more discrete subpopulations differentially distributed along the anteroposterior axis of the spinal cord (Francius et al., 2013, Hayashi et al., 2018). However, specific functions of these IN subsets have not been investigated yet, and the mechanisms that govern their production remain currently unknown.

Recently, we identified Onecut (OC) transcription factors as regulators of neuronal diversification (Kabayiza et al., 2017, Roy et al., 2012, Francius and Clotman, 2014) and of dorsal IN migration (Kabayiza et al., 2017) in the developing spinal cord. OC factors, namely Hepatocyte Nuclear Factor-6 (HNF-6, or OC-1), OC-2 and OC-3, are transcriptional activators present in the digestive tract and in the CNS during embryonic development (Jacquemin et al., 1999, Lemaigre et al., 1996, Jacquemin et al., 2003b, Landry et al., 1997, Vanhorenbeeck et al., 2002). In neural tissue, they regulate production (Espana and Clotman, 2012a), diversification (Roy et al., 2012, Francius and Clotman, 2014, Kabayiza et al., 2017), distribution (Audouard et al., 2013, Espana and Clotman, 2012a, Espana and Clotman, 2012b, Kabayiza et al., 2017) and maintenance (Espana and Clotman, 2012a, Espana and Clotman, 2012b, Stam et al., 2012) of specific neuronal populations, as well as the formation of neuromuscular junctions (Audouard et al., 2012). Here, we demonstrate that OC factors regulate the diversification and the distribution of V2 INs during spinal cord development. Analyzes of OC-deficient embryos showed defective production of specific subpopulations of V2a INs, as well as abnormal distribution of V2a and V2b cells in the developing spinal cord. Furthermore, we uncovered that OC proteins act upstream of specific spinal isoforms of Pou2f2, a POU family transcription factor. Using gain- or loss-of-function experiments, we demonstrated that, as observed for OC factors, Pou2f2 regulates the distribution of V2 INs in the developing spinal cord. Thus, we uncovered a genetic pathway that regulates the diversification and the distribution of V2 INs during embryonic development.

## Results

### OC factors are present in multiple subsets of spinal V2 INs

In the developing spinal cord, OC factors contribute to diversification, migration and maintenance of different neuronal populations (Kabayiza et al., 2017, Roy et al., 2012, Stam et al., 2012). To study V2 IN diversification, we previously established a repertoire of markers that divide embryonic V2 cells into multiple subpopulations (Francius et al., 2013). Although OC factors have been detected in V2 INs (Francius and Clotman, 2010, Francius et al., 2013) and are similarly distributed at distinct antero-posterior levels (Francius et al., 2013), their production in V2 subsets has not been investigated yet. Therefore, we first determined the presence of each OC in these V2 subpopulations at e12.5.

V2a INs include neuronal subsets characterized by the presence of Shox2, MafA, cMaf, Bhlhb5 or Prdm8 (Francius et al., 2013). Only Hnf6 was detected in few Shox2+ V2a cells (Figure 1A-C”; Table 1). In contrast, the 3 OC proteins were detected in MafA+ and cMaf+ V2a subpopulations (Figure 1D-I”; Table 1), but not in Bhlhb5+ or Prdm8+ cells (Table 1; data not shown). V2b INs include similar subsets except for Shox2+ and cMaf+ cells, and contain an additional MafB+ subpopulation (Francius et al., 2013). OC were present in MafA+ but not in MafB+, Bhlhb5+ or Prdm8+ V2b subsets (Figure 1J-L”; Table 1; data not shown). In addition, OC were detected in V2c (non-progenitor Sox1+ cells; Figure 1M-O”; Table 1) but not in V2d (Shox2+Chx10-) cells (Figure 1A-C”; Table 1). Thus, OC factors are present in multiple subpopulations of V2 INs.

**Table 1.** OC factors are present in specific populations and subpopulations of V2 interneurons.

| **V2 – e12.5** |  |  |  |  |
| --- | --- | --- | --- | --- |
| **Populations** | **Subpopulations** | **HNF-6** | **OC-2** | **OC-3** |
| V2a – Chx10 |  | + | + | + |
|  | Shox2 | + | - | - |
|  | MafA | + | + | + |
|  | cMaf | + | + | + |
|  | Bhlhb5 | - | - | - |
|  | Prdm8 | - | - | - |
| V2b – Gata3 |  | + | + | + |
|  | MafA | + | + | + |
|  | MafB | - | - | - |
|  | Bhlhb5 | - | - | - |
|  | Prdm8 | - | - | - |
| V2c – Sox1 |  | + | + | + |
| V2d – Shox2 |  | - | - | - |

The V2 populations, including V2a, V2b, V2c and V2d, are subdivided in smaller subpopulations characterized by differential expression of transcription factors (Francius et al. 2013). OC factors are detected in specific populations and subpopulations of V2 interneurons.

**Figure 1.**
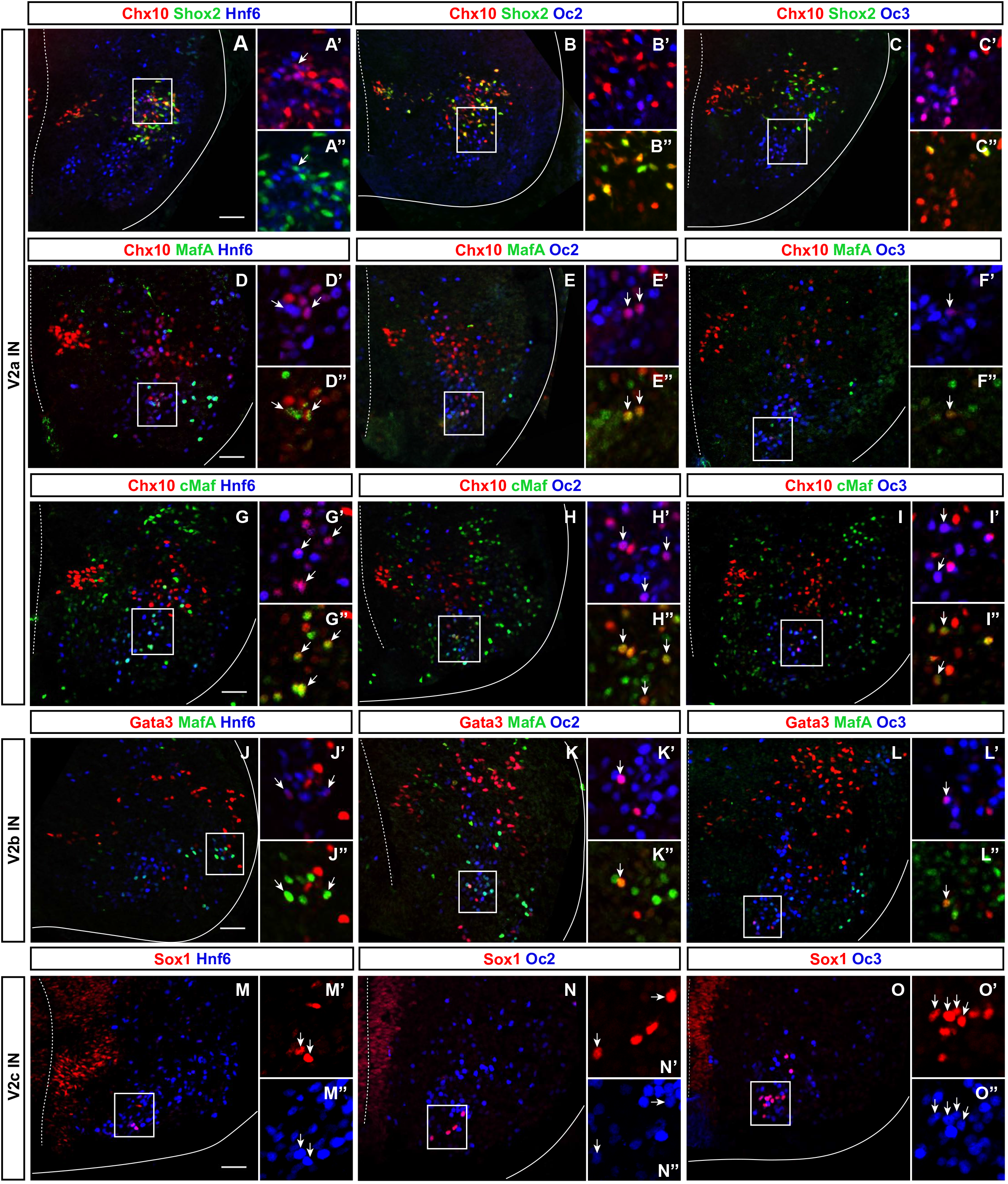
OC factors are present in multiple subsets of V2 interneurons. (A-I”) Immunolabelings for OC, the V2a generic marker Chx10 and markers of V2a subpopulations (Francius et al., 2013) on transverse spinal cord sections (brachial or thoracic levels) of e12.5 wild-type mouse embryos. In each figure, the right ventral quadrant of the spinal cord is shown. Only HNF-6 is detected in Shox2+ V2a cells (arrow in A-C”), whereas the 3 OC are present in the MafA+ and in the cMaf+ V2a subsets (arrows in D-I”). **(J-L”)** Immunolabelings for OC, the V2b generic marker Gata3 and MafA. The 3 OC proteins are detected in MafA+ V2b interneurons (arrows). **(M-O”)** Immunolabelings for OC and the V2c marker Sox1 demonstrate that OC factors are present in a majority of V2c interneurons (arrows). Sox1 in the ventricular zone labels neural progenitors. Scale bar = 50 μm.

### OC factors regulate the diversification of spinal V2 INs

To determine whether OC proteins contribute to the development of V2 IN subsets, we characterized the phenotype of these cells in *Hnf6/Oc2* double-mutant embryos, which lack the three OC factors in the developing spinal cord (Kabayiza et al., 2017, Roy et al., 2012). Given that the number and the distribution of cells in each IN subpopulation vary along the anteroposterior axis of the spinal cord (Francius et al., 2013, Hayashi et al., 2018, Sweeney et al., 2018), this analysis was systematically performed at brachial, thoracic and lumbar levels at e12.5 and e14.5.

In the absence of OC factors, the total number of Chx10+ INs was not significantly changed (Figure 2A-D; Supplementary Figure S1A-B), although a trend toward reduction was detected at brachial level at e12.5 (Figure 2C). Consistently, the number of Chx10+Shox2+ INs was not changed (Figure 2E-H; Supplementary Figure S1C-D”). These observations suggest that OC are not necessary for V2a IN production. In contrast, the smaller V2a subpopulations wherein OC factors were detected in control embryos, characterized by the presence of MafA (Figure 1D-F”) or cMaf (Figure 1G-I”), were almost completely lost in OC mutant embryos (Figure 2I-P; Supplementary Figure S1E-H”). As the total number of Chx10+ and of Shox2+ V2a was not changed (Figure 2A-H; Supplementary Figure S1A-D”), the loss of the MafA+ or cMaf+ subsets may be compensated for by expansion of other V2a subpopulations, markers of which remain to be identified. Nevertheless, our data indicate that OC factors are required either for the expression of V2 subpopulation markers or for the differentiation of specific V2a IN subsets.

**Figure 2.**
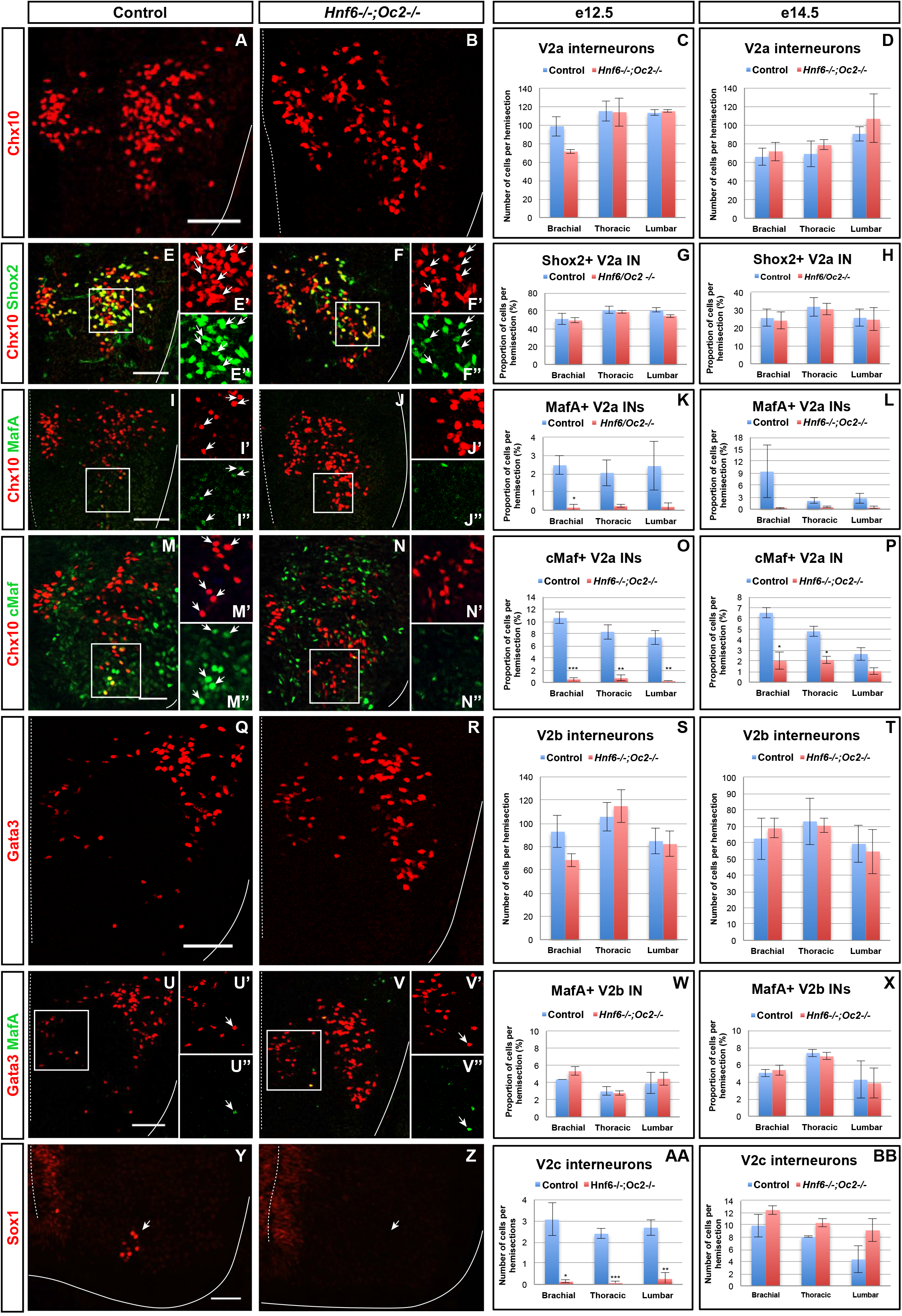
OC factors regulate the diversification of the V2 interneurons. Immunolabelings on transverse spinal cord sections (brachial or thoracic levels) of control or *Hnf6^-/-^;Oc2^-/-^* double-mutant embryos. At e12.5 **(A-C)** and e14.5 **(D)**, the production of the V2a Chx10+ interneurons is not altered in the absence of OC factors. Similarly, the number of Shox2+ V2a is affected neither at e12.5 **(E-G)** nor at e14.5 **(H)**. In contrast, quantitative analysis of control or *Hnf6^-/-^;Oc2^-/-^* littermates at e12.5 **(I-K)** and at e14.5 **(L)** shows reduction in MafA+ V2a interneurons in double mutants as compared to control embryos. Similarly, the number of cMaf+ V2a interneurons is significantly reduced at e12.5 (**M-O**) and e14.5 (**P**) in the absence of OC factors. At e12.5 **(Q-S)** and e14.5 **(T)**, the production of the V2b interneurons is not affected in *Hnf6^-/-^;Oc2^-/-^* embryos. The generation of the MafA+ V2b interneurons is also unchanged at e12.5 **(U-W)** or e14.5 **(X)**. At e12.5 **(Y-AA)**, the number of V2c interneurons is dramatically reduced in the absence of OC factors (Sox1 in the ventricular zone labels neural progenitors). However, this is no longer the case at e14.5 **(BB)**. Mean values ± SEM. * p≤0.05; ** p≤0.01; *** p≤0.001. Scale bar = 50 μm.

To discriminate between these possibilities and to evaluate the contribution of OC factors to V2b diversification, we characterized the phenotype of V2b INs and of their MafA+ subpopulation in the absence of OC proteins. As observed for V2a INs, the total number of V2b cells was not changed in OC mutant embryos (Figure 2Q-T; Supplementary Figure S1I-J), although a trend toward reduction was observed at brachial level at e12.5 (Figure 2S). However, in contrast to V2a, the MafA+ V2b INs were present in normal number in OC mutant embryos (Figure 2U-X; Supplementary Figure S1K-L”). Hence, OC factors are not necessary for the production of the MafA+ V2b subset, although they are required for proper differentiation of the MafA+ and of the cMaf+ V2a subpopulations.

Finally, we assessed the requirement for OC in the production of V2c INs, a V2 population strongly related to V2b cells (Panayi et al., 2010). Although weak production of Sox1 in spinal progenitors was preserved, V2c cells characterized by high Sox1 levels were scarcely detectable in OC mutant embryos at e12.5 (arrows in Figure 2Y-AA). However, the number of V2c was normal at e14.5 (Figure 2BB; Supplementary Figure S1M-N), suggesting that the absence of OC delays the differentiation of V2c INs without affecting the V2b population. Taken together, these observations demonstrate that OC proteins are not required for V2 IN production but regulate the diversification of V2 INs into multiple subpopulations.

### OC factors regulate the distribution of spinal V2 INs

Although the total number of V2a or V2b INs was not affected by the absence of OC factors, careful examination of immunofluorescence labelings suggested that, as observed for spinal dorsal INs (Kabayiza et al., 2017), OC proteins may regulate the distribution of V2 INs in the developing spinal cord (Figure 2A-B; Q-R). Therefore, quantitative distribution analyses (Kabayiza et al., 2017) were performed for each V2 population at brachial, thoracic or lumbar levels at e12.5, namely in the course of ventral IN migration, and at e14.5, i.e. when ventral IN migration in the transverse plane of the spinal cord is completed. Absence of the MafA+ and cMaf+ V2a subpopulations in OC mutants and the small size of other V2 subsets prevented analysis of subpopulation distribution.

At e12.5 in control embryos, V2a INs distributed in 2 connected clusters, a major central group and a minor medial group, at each level of the spinal cord (Figure 3A-C). In mutant embryos, V2a cells similarly distributed in connected central and medial groups. However, the relative cell distribution between the 2 clusters was altered, with less central cells at brachial level and less medial cells at lumbar levels (Figure 3D-L). Altered V2a distribution on the medio-lateral axis was confirmed at e14.5. In control embryos, the 2 V2a groups did coalesce in a more evenly-distributed population that occupied ~70% of the medio-lateral axis (Figure 3M-O). In mutant embryos, V2a INs remained segregated into 2 distinct, although connected, clusters with a majority of cells in medial position (Figure 3P-X). Thus, absence of OC factors perturbs proper distribution of the V2a INs and restricts at e14.5 migration of a fraction of V2a cells in a medial position.

**Figure 3.**
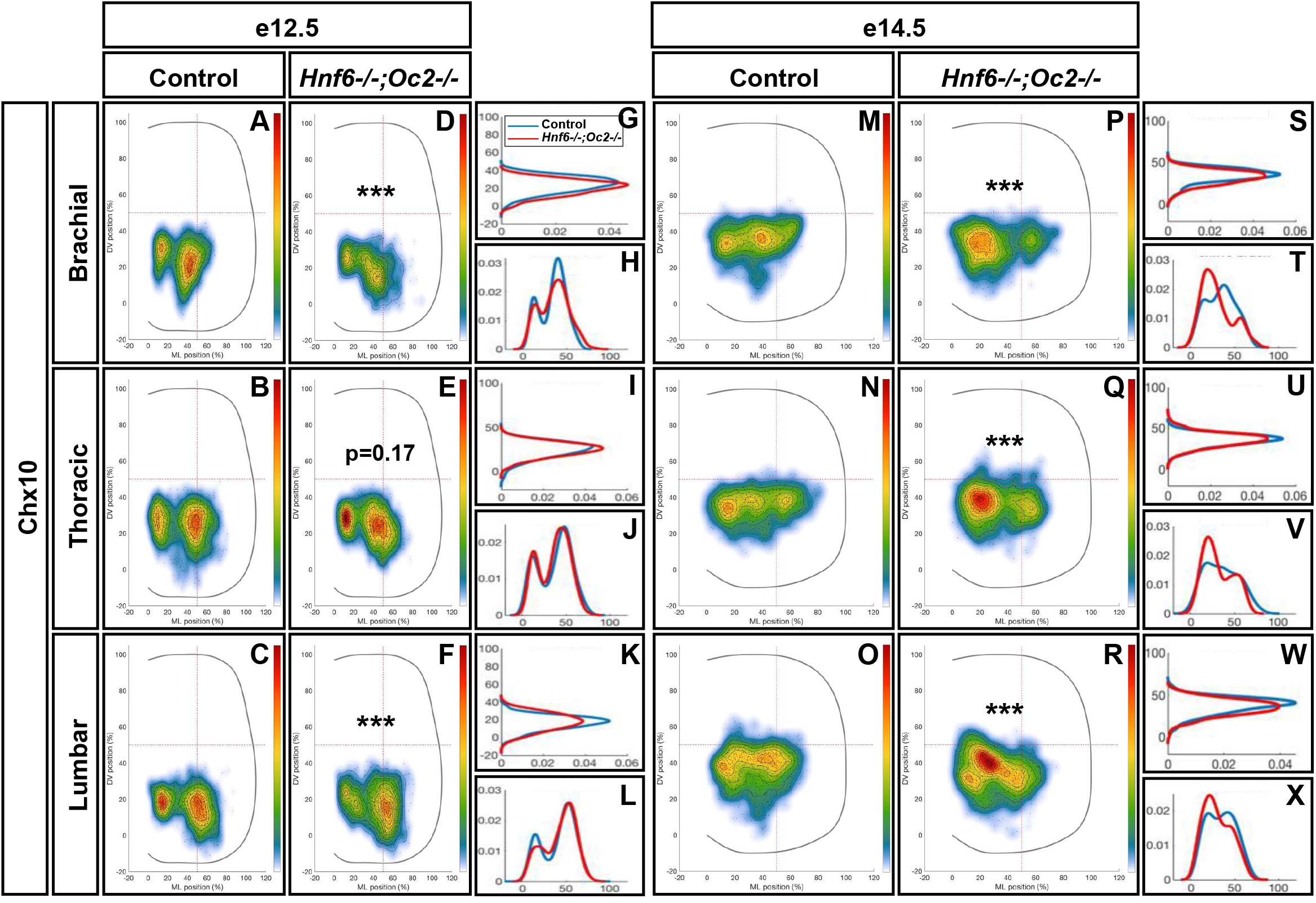
OC factors regulate the distribution of V2a interneurons. Distribution of V2a interneurons on the transverse plane of the spinal cord in control or *Hnf6^-/-^;Oc2^-/-^* doublemutant embryos at brachial, thoracic or lumbar level (only the right hemisection is shown). Two-dimension distribution graphs (left) show integration of cell distribution from multiple sections of multiple embryos of each genotype. One-dimension graphs (right) compare density distribution in control (blue) and in double-mutant embryos (red) on the dorso-ventral (upper) or the medio-lateral (lower) axis of the spinal cord (see Materials and methods for details). **(A-C)** At e12.5 in control embryos, V2a interneurons distribute in 2 connected clusters, a major central group and a minor medial group, at each level of the spinal cord. **(D-L)** In mutant embryos, the relative cell distribution between the 2 clusters seems altered, with relatively less central cells at brachial level and less medial cells at lumbar levels (n=3, p≤0.001 at brachial and lumbar levels; p=0.17 at thoracic level). **(M-X)** Altered V2a distribution on the medio-lateral axis is confirmed at e14.5. **(M-O, S-X)** In control embryos, the 2 V2a groups coalesce in a more evenly-distributed population that occupied ~70% of the medio-lateral axis. **(P-X)** In mutant embryos, V2a interneurons remain segregated into 2 distinct, although connected, clusters with a majority of cells in medial position (n=3, p≤0.001).

To assess whether OC also regulate the position of other V2 populations, we studied the distribution of V2b INs. At e12.5 in control embryos, V2b cells distributed in a major central (brachial level) or lateral (thoracic and lumbar levels) cluster with minor subsets located more medially (arrows in Figure 4A-C) or ventrally (arrowheads in Figure 4A-C). In OC mutant embryos at e12.5, the major population was more compact, more centrally located and slightly more ventral. In addition, the ventral V2b subset was significantly depleted (Figure 4P-X). Consistently, at e14.5, V2b INs in the central cluster remained significantly more compact at thoracic level in the absence of OC factors, and identical trends were observed at brachial and lumbar levels (Figure 4M-X). In addition, a small contingent of V2b migrating towards the medio-dorsal spinal cord in control embryos (arrowheads in Figure 4N,O) was missing in OC mutant littermates (Figure 4Q-X). Taken together, these observations demonstrate that, in addition to V2 diversification, the OC transcription factors regulate proper distribution of V2 INs during spinal cord development.

**Figure 4.**
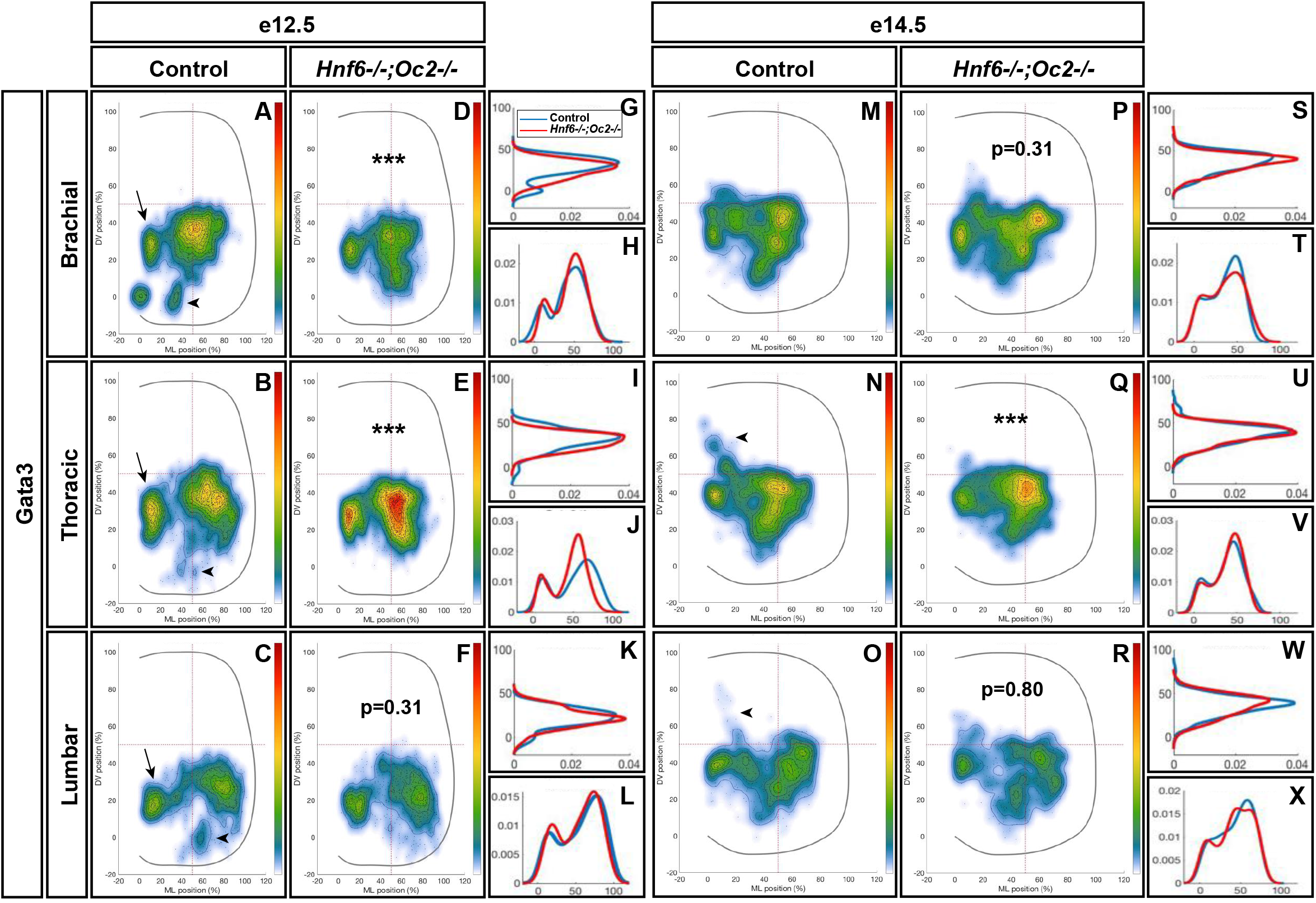
OC factors regulate the distribution of V2b interneurons. **(A-C)** At e12.5 in control embryos, V2b cells are distributed in a major central (brachial level) or lateral (thoracic and lumbar levels) cluster with minor subsets located more medially (arrows) or ventrally (arrowheads). **(D-L)** In OC mutant embryos at e12.5, the major population remains more compact, more centrally located and slightly more ventral. In addition, the ventral V2b subset is significantly depleted (asterisks; n=3, p≤0.001 at brachial and thoracic levels; p=0.31 at brachial level). **(M-X)** Consistently, at e14.5, V2b interneurons in the central cluster are more compact in the absence of OC factors at thoracic level, and a small contingent of V2b migrating towards the medio-dorsal spinal cord in control embryos (arrowheads) is missing in OC mutant littermates (n=3, p≤0.001 at thoracic level; p=0.31 and 0.80 at thoracic and lumbar levels, respectively).

### OC factors control expression of neuronal-specific isoforms of *Pou2f2*

To identify genes downstream of OC that may contribute to V2 IN differentiation and distribution, we performed a microarray comparison of control and of OC-deficient spinal cord transcriptome at e11.5, namely at the stage when significant numbers of V2 cells have been generated and are initiating migration (GEO repository accession number: GSE117871). Among genes showing a differential expression level in the OC mutant spinal cord, *Pou2f2* was significantly upregulated (1.57-fold increase). Pou2f2 (previously named Oct-2) is a transcription factor containing a POU-specific domain and a POU-type homeodomain (Figure 5A) that binds an octamer motif (consensus sequence ATGCAAAT) (Latchman, 1996). *Pou2f2* expression has been detected in B lymphocytes, in neuronal cell lines and in neural tissues including the developing CNS (Lillycrop and Latchman, 1992, Camos et al., 2014, Hatzopoulos et al., 1990). Pou2f2 is required for differentiation of B lymphocytes and for postnatal survival (Corcoran et al., 1993, Corcoran et al., 2004, Hodson et al., 2016, Konig et al., 1995), and is able to modulate neuronal differentiation of ES cells (Theodorou et al., 2009). However, its role in the developing spinal cord remains unknown.

**Figure 5.**
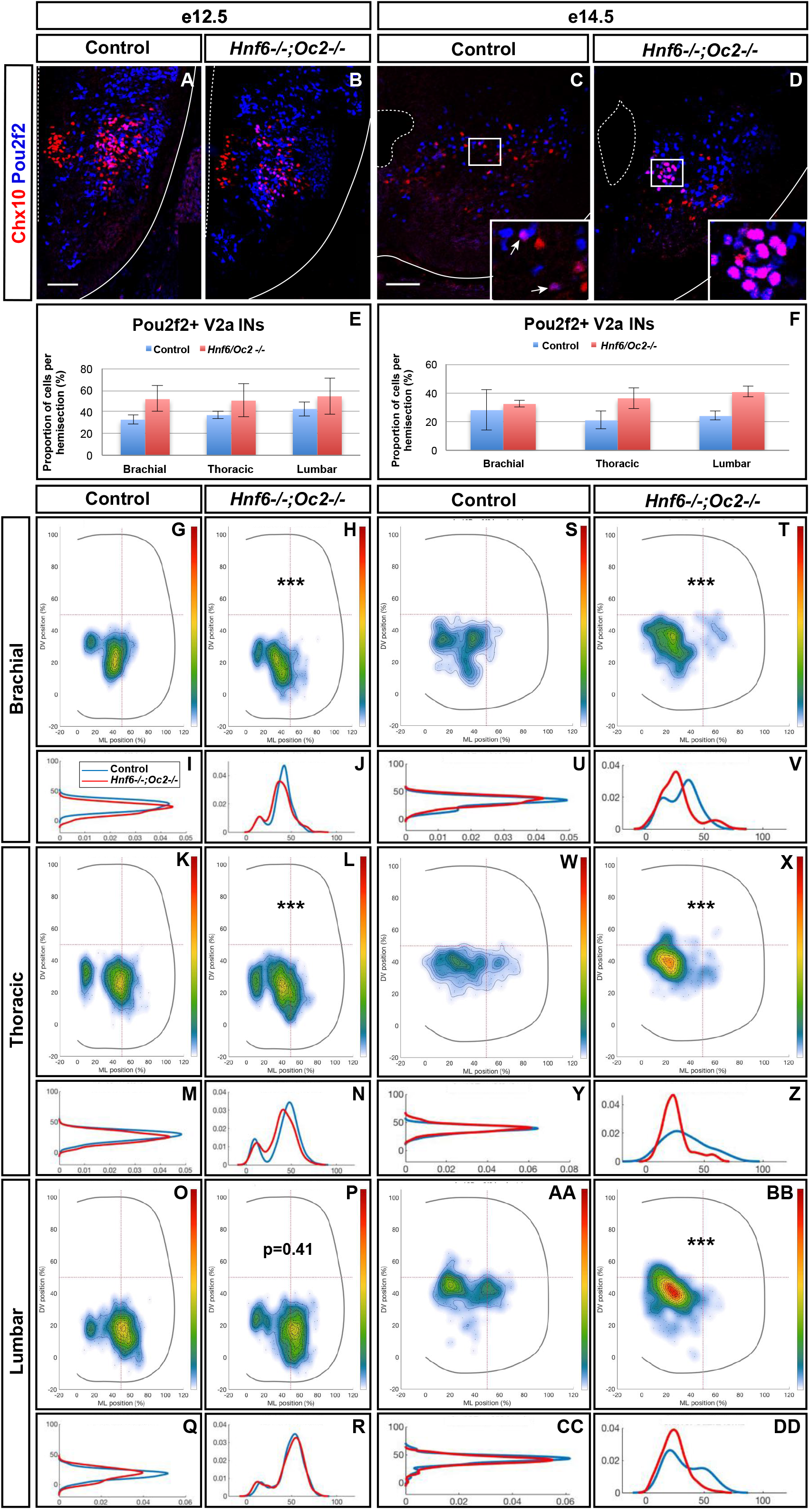
OC factors control expression of spinal cord-specific isoforms of *Pou2f2*. **(A)** The different Pou2f2 isoforms present in the B cells (B_Pou2f2) are characterized by invariant exons (dark grey) and alternative exons 4, 5, 8, 14 or 16 (light grey). They contain a POU-specific domain (light green) encoded by exons 9 and 10 and a POU-type homeodomain (dark green) encoded by exons 11 and 12. The 4 spinal Pou2f2 isoforms (S_Pou2f2.1 to S_Pou2f2.4) (identified in the spinal cord) are characterized by a distinct exon 1 (E1X in light orange), an additional exon E5b (dark orange) and alternative exons E1b and 4 (medium orange and light grey, respectively). The presence of E1b disrupts the reading frame and imposes the use of the ATG located in E2a, whereas the absence of E1b leaves open the use of the ATG located in E1X. The regions corresponding to the generic or to the E5b *in situ* hybridization probes are indicated. **(B-E)** Quantification of spinal Pou2f2 or B-cell isoforms by RT-qPCR. **(B)** In control spinal cords, spinal Pou2f2 isoforms are >30**-**fold more abundant than B-cell isoforms. **(C)** B cell Pou2f2 isoforms barely trend to increase in the absence of OC factors. **(D)** In contrast, spinal Pou2f2 isoforms are 2.6-fold overexpressed in *Hnf6^-/-^;Oc2^-/-^* spinal cords. **(E)** In double mutant spinal cords, spinal Pou2f2 isoforms are >60-fold more abundant than B-cell isoforms. **(F-I)** *In situ* hybridization labelings on transverse sections (brachial level) of control or *Hnf6^-/-^;Oc2^-/-^* spinal cords at e11.5 with **(F-G)** a generic *Pou2f2* probe complementary to spinal and to B-cell isoforms **(A)** or **(H-I)** a spinal isoform-specific probe corresponding only to exon E5b **(A)**. **(F, H)** In control embryos, *Pou2f2* is strongly expressed in ventral and in dorsal interneuron populations, and more weakly in the ventral motor neuron area. **(G, I)** In OC mutant embryos, *Pou2f2* is upregulated in interneuron populations and its expression is expanded in ventral populations (arrowheads) and in the motor neurons (arrows). * p≤0.05; ** p≤0.01. Scale bars = 50 μm.

Based on work in B cells, multiple *Pou2f2* isoforms generated by alternative splicing have been described (Figure 5A; (Lillycrop and Latchman, 1992, Wirth et al., 1991, Hatzopoulos et al., 1990, Liu et al., 1995, Stoykova et al., 1992)). Therefore, we first determined whether similar isoforms are found in the developing spinal cord. However, we systematically failed to obtain RT-PCR products using upstream primers in described exon 1 (asterisks in Supplementary Figure S2A-B and data not shown; Table 2) and amplifications encompassing exons 5 to 6 generated predominant amplicons larger than expected (arrowheads in Supplementary Figure S2B; Table 2), suggesting the existence of alternative exons in *Pou2f2* embryonic spinal cord transcripts (Figure 5A). Data mining the NCBI Nucleotide database for *Pou2f2* sequences identified a predicted murine *Pou2f2* isoform (X6 sequence, accession number XM_006539651.3) with a different exon 1 (E1X) and an additional sequence between exons 5 and 6, the size of which (279 bp) corresponded to the size differences estimated in our amplifications encompassing exons 5 to 6 (arrowheads in Supplementary Figure S2B; Table 2). Using PCR primers in this predicted sequence, we were able to amplify a 5’ region of *Pou2f2* from the alternative E1X exon and an additional sequence between exons 5 and 6 (Supplementary Figure S2C; Table 2), suggesting that alternative isoforms similar to this predicted sequence are produced in the developing spinal cord. However, amplifications from E1X systematically produced 2 amplicons (arrowheads in Supplementary Figure S2C; Table 2), suggesting the existence of an alternative exon downstream to E1X. Sequencing of our PCR products and alignment to genomic DNA confirmed that predominant *Pou2f2* isoforms in the developing spinal cord contain the alternative E1X exon, an additional exon (E5b) between exons 5 and 6, and can undergo alternative splicing of a short (61bp) exon (E1b) between E1X and exon 2 (Supplementary Figure S2D). E5b exon maintains the reading frame. In contrast E1b exon disrupts it, imposing the use of the ATG located in exon 2 to generate a functional Pou2f2 protein, whereas the absence of E1b leaves open the use of an alternative upstream ATG located at the 3′ end of E1X (Figure 5A; Supplementary Figure S2D). Hence, our RT-PCR and sequencing data indicate that 4 neuronal *Pou2f2* isoforms different from the previously described B-cell or neural isoforms are produced in the developing spinal cord (Figure 5A).

**Table 2.** Pou2f2 isoforms in the developing spinal cord are different from B-cell isoforms.

| **Amplification** | **Expected sizes** | **B cells** | **Control spinal cord** |
| --- | --- | --- | --- |
| **E1 → E6** | 272 bp<br>390 bp<br>456 bp | 272 bp<br>390 bp<br>456 bp | No amplification |
| **E7 → E9** | 174 bp<br>222 bp | 174 bp<br>222 bp | 174 bp |
| **E10 → E12** | 452 bp | 452 bp | 452 bp |
| **E13 → E17** | 160 bp<br>296 bp<br>370 bp | 160 bp<br>296 bp<br>370 bp | 370 bp |
| **E1 → E7** | 348bp<br>466 bp<br>532 bp | 466 bp<br>532 bp | No amplification |
| **E2 → E7** | 284 bp<br>402 bp<br>468 bp | 402 bp<br>468 bp | ~ 680 bp<br>~ 750 bp |
| **E3 → E6** | 154 bp<br>272 bp<br>378 bp | 154 bp<br>272 bp<br>378 bp | 272 bp<br>378 bp<br>~ 550 bp<br>~ 650 bp |
| **E4 → E7** | 340 bp | 340 bp | ~ 600 bp |
| **E5 → E7** | 201 bp | 201 bp | 201 bp<br>~ 480 bp |
| **E6 → E7** | 160 bp | 160 bp | 160 bp |
| **E1X → E3** | 152 bp | No amplification | 152 bp<br>~ 220 bp |
| **E1X → E5b** | 554 bp<br>620 bp | N.D | 554 bp<br>620 bp |
| **E5b** | 234 bp | N.D | 234 bp |
| **E3 → E5b** | 429 bp<br>495 bp | N.D | 429 bp<br>495 bp |
| **E5b → E7** | 416 bp | N.D | 416 bp |

Regions covering B lymphocytes or predicted Pou2f2 isoforms (exons E1 or E1X to E17) were amplified by RT-PCR from B lymphocyte or embryonic spinal cord RNA.
Orange cells = unexpected results.

However, minor transcripts corresponding to B-cell isoforms are also detected in the embryonic spinal cord (Supplementary Figure S2A-B; Table 2). To assess the relative abundance of each transcript type in this tissue and to evaluate the extent of their relative overexpression in the absence of OC factors, we quantified each isoform type in control and in OC-deficient spinal cord at e11.5. In control spinal cords, spinal *Pou2f2* isoforms were >30-fold more abundant than B-cell isoforms (Figure 5B), consistent with our RT-PCR observations (Supplementary Figure S2). In the absence of OC factors, spinal isoforms were ~2.6-fold overexpressed whereas B-cell isoforms barely trended to increase (Figure 5C-E). Thus, OC factors repress expression of spinal *Pou2f2* isoforms in the developing spinal cord. To confirm these data and to determine the expression pattern of *Pou2f2* in the ventral spinal cord, *in situ* hybridization was performed on sections from control or *Hnf6/Oc2* double-mutant spinal cords using either a generic *Pou2f2* probe complementary to spinal and to B-cell isoforms or a spinal isoform-specific probe corresponding only to exon E5b (Figure 5A). Using the generic *Pou2f2* probe on control tissue, we detected *Pou2f2* transcripts in the ventral region of the spinal cord, with lower expression levels in the location of the motor columns (arrows in Figure 5F). In OC mutant embryos, *Pou2f2* expression was globally increased and additionally expanded in the ventral area (arrowheads in Figure 5F-G) including the motor neuron territories (arrows in Figure 5F-G). Similar observations were made with the spinal isoform-specific probe (Figure 5H-I). Thus, OC factors restrict and moderate *Pou2f2* expression in ventral spinal populations likely including V2 INs.

### Pou2f2-positive V2 INs are mislocated in the absence of OC factors

To assess whether increased *Pou2f2* expression in *Hnf6^-/-^;Oc2^-/-^* spinal cords corresponded to an expansion of Pou2f2 distribution in V2 INs or an upregulation in its endogenous expression territory, we first quantified the number and distribution of Pou2f2-containing Chx10+ cells at e12.5 and e14.5 (Figure 6; Supplementary Figure S3). Immunofluorescence experiments demonstrated that Pou2f2 is present in V2a INs in control embryos (Figure 6A,C; Supplementary Figure S3A,D), although it was only sparsely detected in MafA+ and cMaf+ V2a subsets (data not sown). Intensity of the labeling confirmed increased Pou2f2 production in the ventral regions of the spinal cord in OC mutant embryos (Figure 6A-F). However, the number in Pou2f2-positive V2a INs was not significantly different (Figure 6E-F; Supplementary Figure S3), suggesting that Pou2f2 is increased in its endogenous expression domain. In contrast, the distribution of Pou2f2-positive V2a INs was affected (Figure 6A-D, 6G-DD). At e12.5, cells in the central clusters were slightly reduced at brachial and at thoracic levels (Figure 6G-R). In contrast, at e14.5, a majority of V2a containing Pou2f2 settled in a medial position (Figure 6S-DD), reminiscent of the distribution defects observed for the whole V2a population (Figure 3). Similarly, Pou2f2 was detected in V2b INs in control embryos, although in a more restricted number of cells (Figure 7A-B, E). In the absence of OC factors, the number of Pou2f2-positive V2b cells was not significantly increased (Figure 7E-F) but, as observed for V2a, this subset of V2b INs was mislocated with cells more central at e12.5 and more clustered on the medio-lateral axis at e14.5 (Figure 7G-DD). Taken together, these observations suggest that Pou2f2 may contribute to control V2 IN migration during spinal cord development and could participate in alterations of V2 distribution in the absence of OC factors.

**Figure 6.**
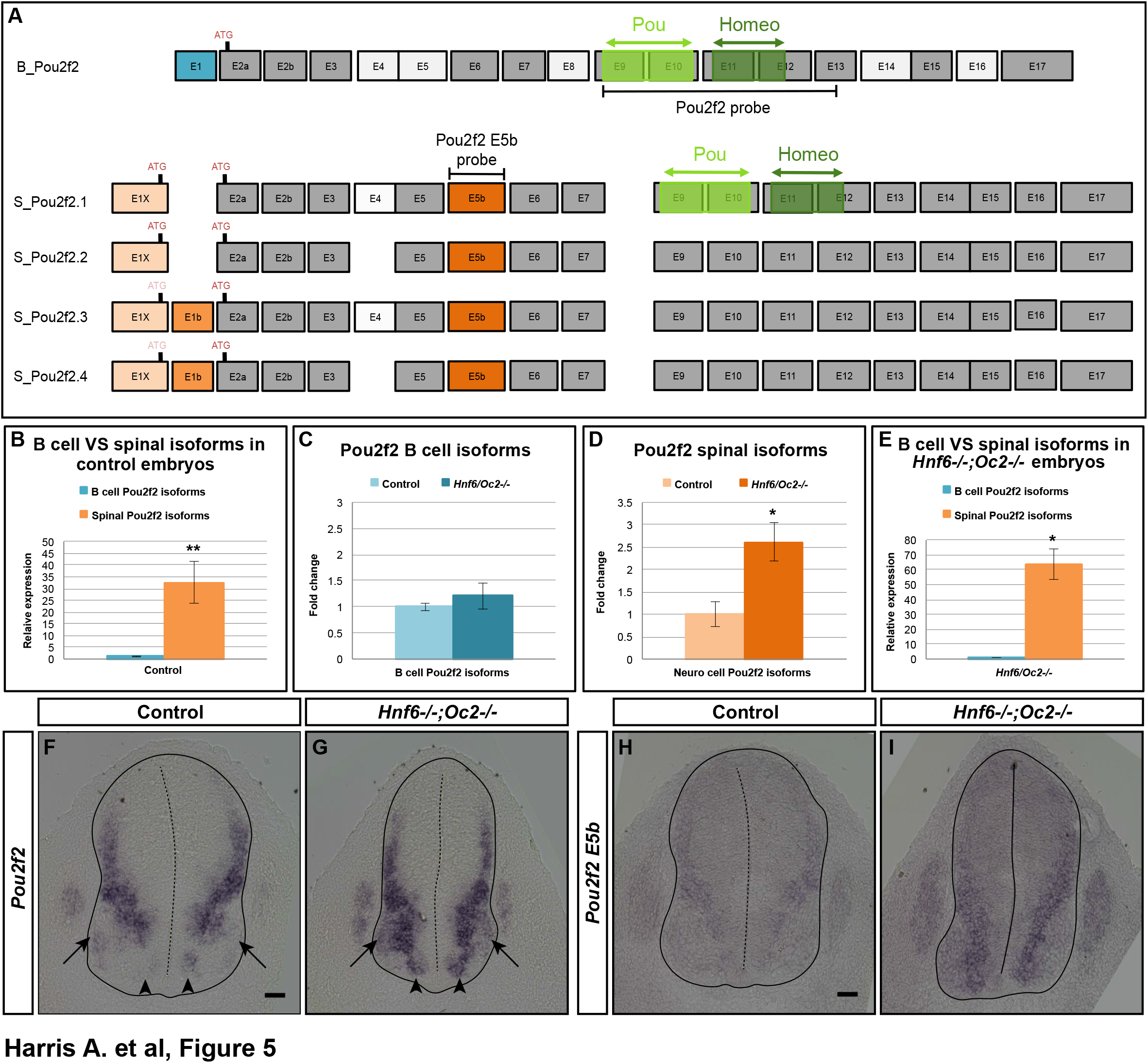
The Pou2f2+ V2a interneurons are mislocated in the absence of OC factors. **(A-F)** Immunolabelings and quantification of Pou2f2+ V2a interneurons in control or *Hnf6^-/-^;Oc2^-/-^* mutant embryos. At e12.5 **(A-B)** and e14.5 **(C-D)**, Pou2f2 is detected in V2a Chx10^+^ interneurons, and the number of Pou2f2-containing Chx10+ cells trends to increase but is not significantly different in the absence of OC factors **(E-F)**. **(G-DD)** Distribution of Pou2f2+ V2a interneurons on the transverse plane of the spinal cord in control or *Hnf6^-/-^;Oc2^-/-^* double-mutant embryos. One-dimension graphs (lower) show density distribution on the dorso-ventral (left) or the medio-lateral (right) axis of the spinal cord. **(G-R)** At e12.5, cells in the central clusters are slightly reduced at brachial and at thoracic levels in the absence of OC factors (n=3, p≤0.001 at brachial and thoracic levels; p=0.41 at lumbar level). **(S-DD)** At e14.5, a vast majority of V2a containing Pou2f2 settles in a more medial position in *Hnf6^-/-^;Oc2^-/-^* spinal cords (n=3, p≤0.001). Mean values ± SEM. Scale bar = 50 μm.

**Figure 7.**
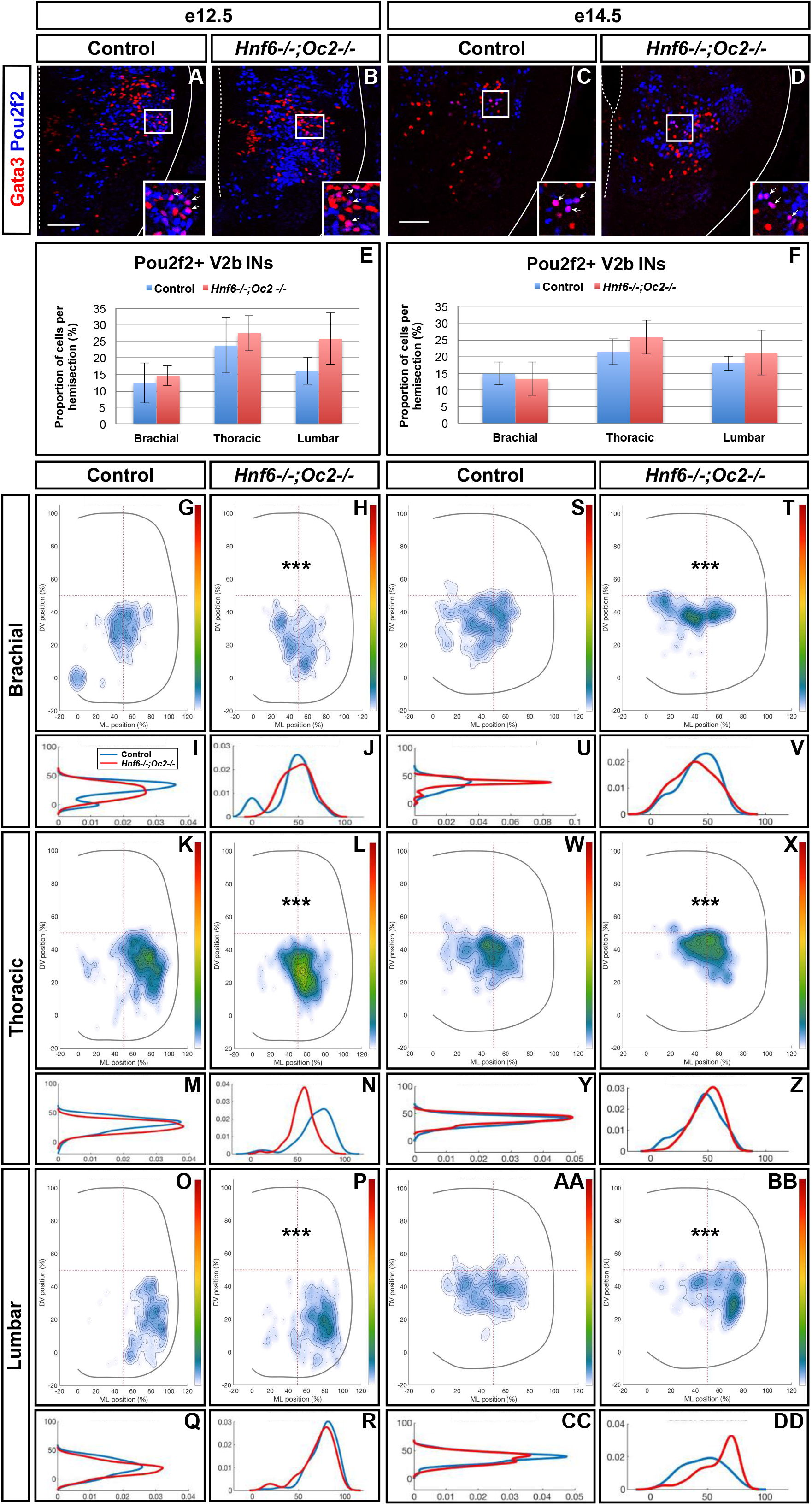
The Pou2f2+ V2b interneurons are mislocated in the absence of OC factors. **(A-F)** Immunolabelings and quantification of Pou2f2+ V2b interneurons in control or *Hnf6^-/-^;Oc2^-/-^* mutant embryos. At e12.5 **(A-B)** and e14.5 **(C-D)**, Pou2f2 is present in V2b Gata3+ interneurons, but the number of Pou2f2+ V2b cells was not significantly increased in the absence of OC factors **(E-F)**. **(G-DD)** Distribution of Pou2f2+ V2b interneurons on the transverse plane of the spinal cord in control or *Hnf6^-/-^;Oc2^-/-^* double-mutant embryos. One-dimension graphs (lower) show density distribution on the dorso-ventral (left) or the medio-lateral (right) axis of the spinal cord. **(G-R)** At e12.5, Pou2f2-containing V2b are more central and slightly more ventral in the absence of OC factors (n=3, p≤0.001). **(S-DD)** At e14.5, this subset of V2b interneurons is more clustered on the medio-lateral axis in *Hnf6^-/-^;Oc2^-/-^* spinal cords (n=3, p≤0.001). Mean values ± SEM. Scale bar = 50 μm.

### Pou2f2 regulates the distribution of spinal V2 INs

To determine whether Pou2f2 is able to modulate V2 IN distribution, we overexpressed *Pou2f2* in the chicken embryonic spinal cord (Supplementary Figure S4). Increased Pou2f2 did not impact on the number of V2a or V2b (Figure 8A-F). In contrast, it did alter V2 IN location. In HH27-28 control spinal cord, V2a INs were distributed in 2 closely connected clusters on the medio-lateral axis of the neuroepithelium (Figure 8G). In electroporated spinal cord, lateral migration was increased and a majority of V2a INs were clustered in a single group in a central position (Figure 8H-J) with ectopic lateral extensions (arrow in Figure 8H). In control spinal cord, V2b were distributed in 2 groups along the medio-lateral axis with a majority of cells in the lateral cluster (Figure 8K). In electroporated spinal cord, the V2b INs were equally distributed between these 2 clusters (Figure 8L-N). Thus, consistent with our observation in OC mutant embryos, increased Pou2f2 can modulate the distribution of V2 INs in the developing spinal cord.

**Figure 8.**
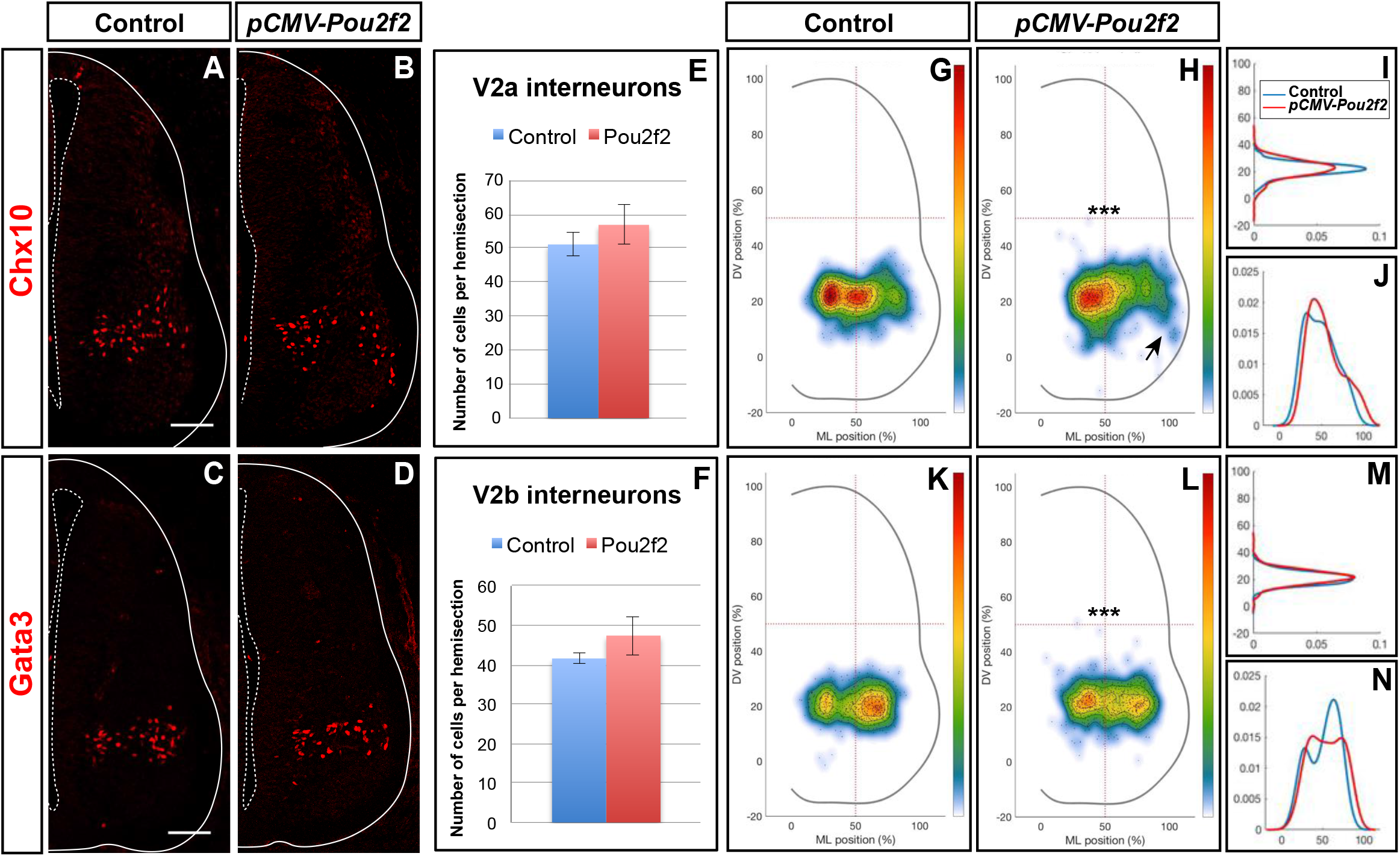
V2 interneuron distribution is altered after misexpression of *Pou2f2*. Overexpression of *Pou2f2* in chick embryonic spinal cord after electroporation at HH14-16 and immunolabelings 72 hours after electroporation. **(A-F)** At HH27-28, *Pou2f2* overexpression does not impact the number of V2a **(E)** or V2b **(F)** interneurons. In contrast, it alters V2 distribution. **(G-J)** In control spinal cord, V2a interneurons are distributed in two closely connected clusters on the medio-lateral axis of the neuroepithelium. In electroporated spinal cord, lateral migration is increased and a majority of V2a interneurons are clustered in a single central group with ectopic lateral extensions (arrows; n=3, p≤0.001). **(K-N)** In control spinal cord, V2b are distributed in two groups along the medio-lateral axis with a majority of cells in the lateral cluster. In electroporated spinal cord, the V2b interneurons are equally distributed between these 2 clusters (n=3, p≤0.001). Mean values ± SEM. Scale bar = 50 μm.

To confirm the influence of Pou2f2 on V2 migration, we studied V2 distribution in mouse embryos devoid of Pou2f2 (Corcoran et al., 1993) at e12.5. Absence of Pou2f2 did not impact on the number of V2a INs (Figure 9A-C) nor on the Shox2+, cMaf+ or MafA+ V2a subsets (Supplementary Figure S5). In contrast, V2a distribution was affected in *Pou2f2* mutants. As compared to the two V2a clusters observed in control embryos, Chx10+ cells were more scattered in *Pou2f2* mutant embryos (Figure 9D-O). Furthermore, although the number of V2b, V2c or MafA+ V2b cells was not changed in the absence of Pou2f2 (Figure 10A-C; Supplementary Figure S6), V2b cells remained more medial at brachial and thoracic levels and segregated more extensively on the medio-lateral axis at lumbar levels (Figure 10D-O). Taken together, these observations demonstrate that Pou2f2 regulate the distribution of V2 INs during spinal cord development.

**Figure 9.**
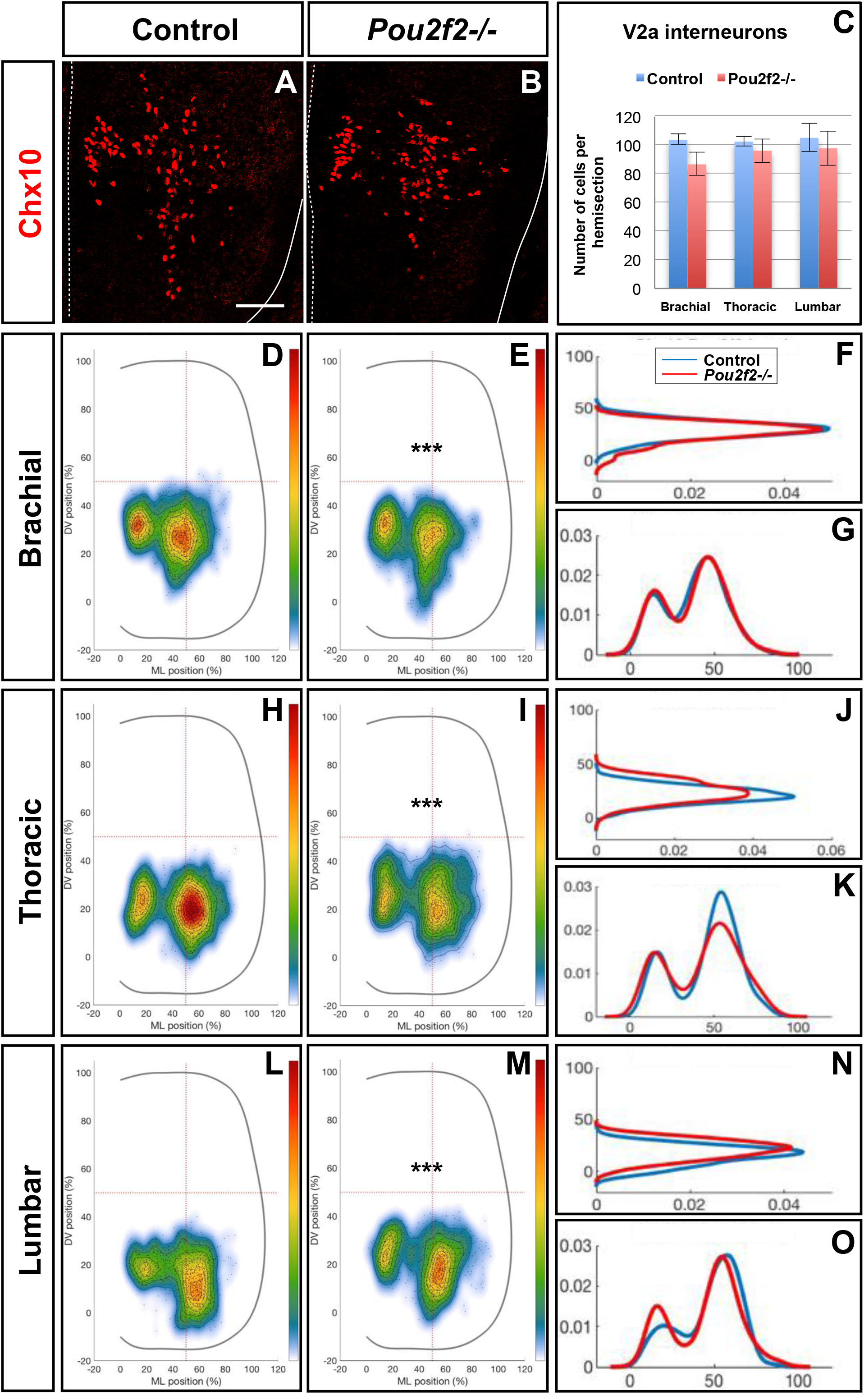
Pou2f2 regulate the distribution of V2a interneurons. **(A-C)** Immunolabelings and quantification of V2a interneurons in control or *Pou2f2^-/-^* mutant embryos. At e12.5, the production of the Chx10+ V2a interneurons is not altered in absence of Pou2f2. **(D-O)** Distribution of V2a and Shox2+ V2a interneurons on the transverse plane of the spinal cord in control or *Pou2f2^-/-^* mutant embryos. One-dimension graphs (right) show density distribution on the dorso-ventral (upper) or the medio-lateral (lower) axis of the spinal cord. V2a distribution is affected in *Pou2f2^-/-^* mutants. As compared to the two V2a clusters observed in control embryos, Chx10+ cells are relatively more abundant in the medial cluster in *Pou2f2^-/-^* mutant embryos (n=3, p≤0.001). Mean values ± SEM. Scale bar = 50 μm.

**Figure 10.**
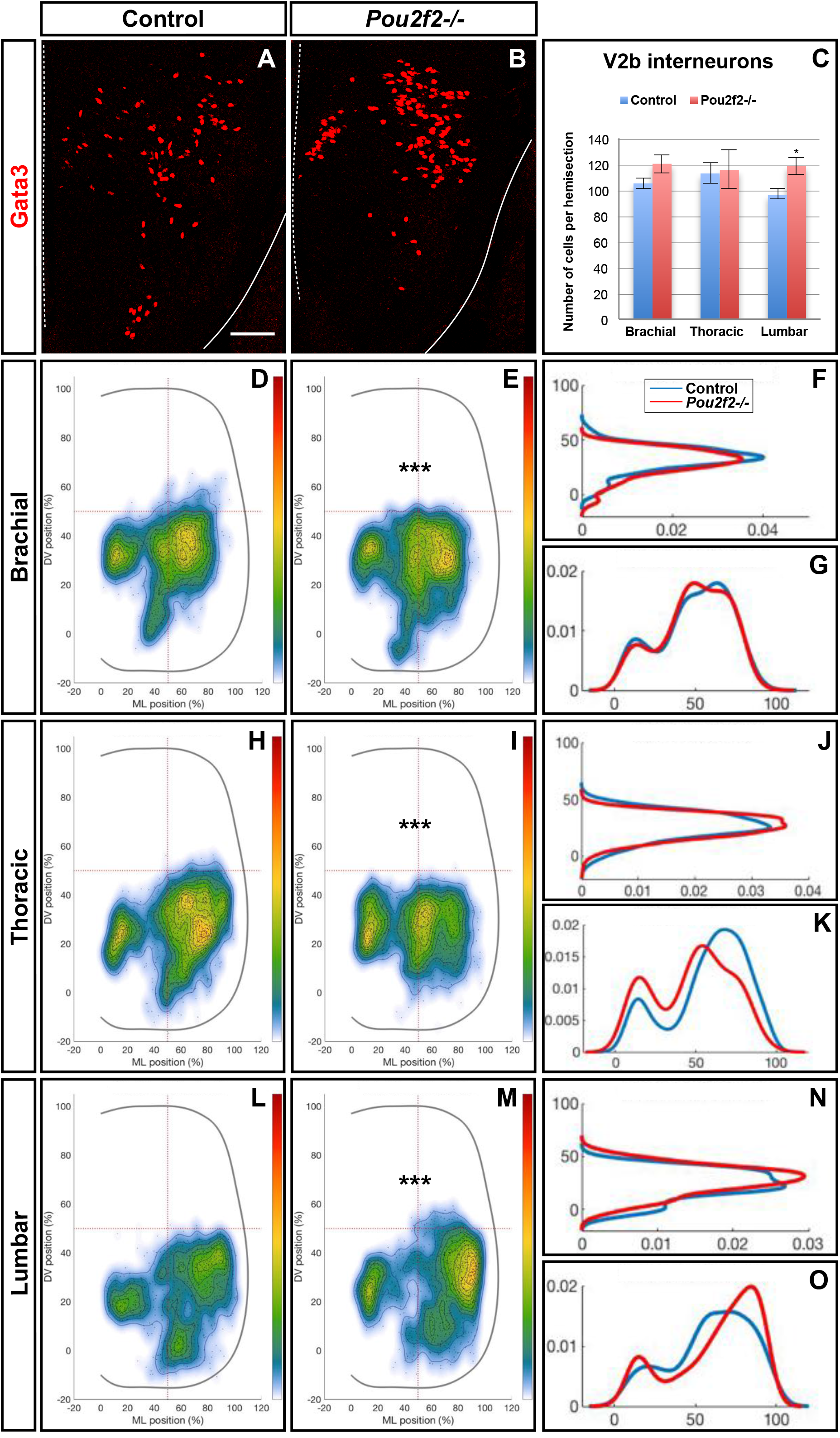
Pou2f2 regulate the distribution of V2b interneurons. **(A-C)** Immunolabelings and quantification of V2b interneurons in control or *Pou2f2^-/-^* mutant embryos. At e12.5, the production of the Gata3+ V2b interneurons is not affected by the absence of Pou2f2. **(D-O)** Distribution of V2b interneurons on the transverse plane of the spinal cord in control or *Pou2f2^-/-^* mutant embryos. One-dimension graphs (right) show density distribution on the dorso-ventral (upper) or the medio-lateral (lower) axis of the spinal cord. The distribution of V2b cells is altered in *Pou2f2^-/-^* mutants, as V2b interneurons remained more medial at thoracic level and segregated more extensively on the medio-lateral axis at lumbar level (n=3, p≤0.001). Mean values ± SEM. Scale bar = 50 μm.

## Discussion

In the recent years, several studies demonstrated that proper distribution of neuronal populations and subpopulations in the developing spinal cord is critical for adequate formation of spinal circuits (Bikoff et al., 2016, Goetz et al., 2015, Hayashi et al., 2018, Hilde et al., 2016, Surmeli et al., 2011, Tripodi et al., 2011). However, the genetic programs that control the diversification of spinal neuronal populations into specialized subpopulations and the proper settling of these neuronal subsets in the spinal parenchyma remain elusive. Here, we provide evidence that OC transcription factors regulate the diversification of spinal V2 INs, and that a genetic cascade involving OC factors and their downstream target Pou2f2 controls the distribution of V2 INs in the developing spinal cord.

### Control of V2 IN diversification by the OC factors

Cardinal populations of spinal ventral INs have been well characterized, and their global contribution to the activity of motor circuits has been extensively studied (reviewed in (Boije and Kullander, 2018, Gosgnach et al., 2017, Ziskind-Conhaim and Hochman, 2017). However, more recently, the idea emerged that these cardinal populations are not homogeneous ensembles but rather contain multiple neuronal subsets with distinct molecular identities and functional properties (Bikoff et al., 2016, Borowska et al., 2015, Borowska et al., 2013, Francius et al., 2013, Sweeney et al., 2018, Talpalar et al., 2013). V2a INs comprise two major divisions, namely type I and type II V2a cells, that are arrayed in counter-gradients along the antero-posterior axis of the spinal cord and activate different patterns of motor output at brachial or lumbar levels. Furthermore, these two large divisions can themselves be fractionated at birth into 11 subsets characterized by distinct combinations of markers, differential segmental localization and specific distribution patterns on the medio-lateral axis of the spinal cord (Hayashi et al., 2018). In the zebrafish, 3 distinct subclasses of V2a INs participate in separate microcircuit modules driving slow, intermediate or fast motor neuron activity (Ampatzis et al., 2014). Taken together, these observations suggest that cardinal IN populations only constitute the first organization level of functionally distinct neuronal subsets that contribute to diversity and flexibility within spinal motor circuits.

We showed here that OC factors are present in subsets of V2 INs and contribute to their diversification. Normal numbers of cardinal V2a and V2b cells were generated in OC mutant embryos, suggesting that these factors do not contribute to the production of V2 cells (Clovis et al., 2016, Lee et al., 2008, Thaler et al., 2002) nor to the segregation of the V2a and V2b lineages through differential activation of Notch signaling (Del Barrio et al., 2007, Joshi et al., 2009, Misra et al., 2014, Peng et al., 2007). In contrast, V2a subpopulations characterized by the presence of MafA or cMaf were strongly depleted in the absence of OC proteins. This observation results either from a loss of these V2a subsets or from a downregulation of *MafA* or *cMaf* expression in these cells. Uncomplete knowledge of the whole collection of V2a subsets prevented to assess whether this apparent loss of specific subpopulations was compensated for by an expansion of neighboring subsets. Nevertheless, these data demonstrate altered differentiation of V2 IN subsets in the absence of OC factors, as previously observed for spinal motor neurons (Roy et al., 2012) and dorsal INs (Kabayiza et al., 2017). In addition, the production of V2c cells was delayed in OC mutant embryos, although V2b that are supposed to constitute the source of V2c (Panayi et al., 2010) were timely generated. This points to a specific contribution of OC protein to the development of V2c INs, the mechanism of which is currently unknown.

### Control of V2 IN distribution by the OC factors

Beside diversification, the characterization of functionally distinct IN subpopulations unveiled a strong correlation between the distribution of each IN subset and their contribution to distinct microcircuit modules (Bikoff et al., 2016, Borowska et al., 2013, Goetz et al., 2015, Hayashi et al., 2018, Hilde et al., 2016, Tripodi et al., 2011). These data support a model wherein correct localization of spinal IN subsets is critical for proper formation of sensory and sensory-motor circuits, highlighting the importance of a strict regulation of short-distance neuronal migration in the developing spinal cord. However, genetic determinants that control spinal IN migration have only been sparsely identified. Sim1 regulate ventro-dorsal migration of the V3 IN subsets (Blacklaws et al., 2015). Similarly, SATB2 control the position of inhibitory sensory relay INs along the medio-lateral axis of the spinal cord (Hilde et al., 2016). Here, we provide evidence that the OC factors control a genetic program that regulates proper positioning of V2 INs during embryonic development. In the absence of OC proteins, a fraction of V2a INs remained in a more medial location, expanding the medial cluster containing locally-projecting cells at the expanse of the lateral cluster that comprises the supraspinal-projecting V2a INs (Hayashi et al., 2018). V2b alterations were less spectacular, although ventral and dorsal contingents were reduced and the cell distribution in the central cluster was altered. Variability in the alterations observed at e12.5 and e14.5 highlights that spinal migration is not completed at e12.5 and suggests that distribution of earlier- or later-migrating neurons may be differently affected by the absence of OC proteins. In addition, migration along the anteroposterior axis, which can not be analyzed in our experimental setup, could also be perturbed. Alterations of the distribution of V2a and V2b interneurons likely correlate with alterations in their differentiation program. Indeed, the production of adequate clues to ensure proper cell positioning is an intrinsic component of any neuronal differentiation program (Blacklaws et al., 2015, Hayashi et al., 2018, Hilde et al., 2016). In any case, our observations are consistent with the contribution of OC factors to the migration of several populations of spinal dorsal INs (Kabayiza et al., 2017). This raises the question whether similar cues might be regulated by identical genetic programs and used by different IN populations to organize proper distribution of ventral and dorsal IN subsets in the developing spinal cord. Identification of the factors downstream of OC protein involved in the control of neuronal migration will be necessary to answer this question.

### An OC-Pou2f2 genetic cascade regulates the migration of V2 INs

Therefore, we attempted to identify genes downstream of OC factors and possibly involved in the control of IN migration using a global approach comparing the transcriptome of whole spinal cords isolated from control or OC-deficient embryos. Absence of known regulators of neuronal migration among the most affected genes suggests that different actors may be active downstream of OC proteins in distinct IN populations to regulate proper neuronal distribution. In contrast, we uncovered that expression of the transcription factor *Pou2f2* is repressed by OC factors in different spinal populations. Surprisingly, our data demonstrated that variant *Pou2f2* isoforms are produced in the developing spinal cord as compared to B lymphocytes (Lillycrop and Latchman, 1992, Wirth et al., 1991, Hatzopoulos et al., 1990, Liu et al., 1995, Stoykova et al., 1992), and that these spinal variants are regulated by OC proteins. Spinal-enriched transcripts encode Pou2f2 proteins containing additional peptidic sequences upstream of the POU-specific domain and of the homeodomain. Furthermore, exon 1 is different and corresponds to sequences located ~47kb upstream of the transcription initiation site used in B cells (data not shown) in the mouse genome, suggesting that OC regulate *Pou2f2* expression from an alternative promoter. However, we cannot exclude that additional exon(s) could be present upstream of the identified sequences, and determination of the regulating sequences targeted by the OC protein will require thorough characterization of the produced transcripts. In addition, we cannot rule out indirect regulation of *Pou2f2* expression by the OC factors, as OC are usually considered to be transcriptional activators rather than repressors (Beaudry et al., 2006, Jacquemin et al., 2000, Jacquemin et al., 2003a, Lannoy et al., 2000, Pierreux et al., 2004, Roy et al., 2012).

Nevertheless, our observations demonstrate that Pou2f2 is downstream of OC factors in the V2 INs and also contributes to regulate the distribution of V2 INs during embryonic development. The number of Pou2f2-containing V2 was not significantly increased in OC mutant spinal cords, suggesting that the absence of OC protein resulted in relaxing of Pou2f2 production in its endogenous expression domain rather that ectopic activation in other V2 subsets. Increased production of Pou2f2 in the chicken embryonic spinal cord resulted in alterations in the localization of V2 populations without any change in cell number, pointing to a possible contribution of Pou2f2 to the regulation of V2 migration downstream of OC factors. Consistently, V2 distribution was perturbed in *Pou2f2* mutant embryos without any alteration in V2 population or subpopulation cell numbers, demonstrating the involvement of Pou2f2 in the control of V2 IN distribution. Alterations in V2 distribution after *Pou2f2* electroporation were not comparable to that observed in OC mutant embryos and were not strictly opposite to *Pou2f2* knockout phenotype because the developmental stages obtained after chicken embryo electroporation were much earlier than the developmental stages of analyzed mouse embryos. Nevertheless, our data demonstrate that a genetic cascade comprising OC and Pou2f2 transcription factors ensures proper distribution of V2 cells during spinal cord development. This program may not be restricted to V2 cells, as diversification and distribution of dorsal INs and of motor neurons are also altered in the absence of OC factors (Kabayiza et al., 2017, Roy et al., 2012) and as *Pou2f2* expression in the OC mutant spinal cord is increased in multiple neuronal populations.

## Materials and methods

### Ethics statement and mouse lines

All experiments were strictly performed in accordance with the European Community Council directive of 24 November 1986 (86-609/ECC) and the decree of 20 October 1987 (87-848/EEC). Mice were raised in our animal facilities and treated according to the principles of laboratory animal care, and experiments and mouse housing were approved by the Animal Welfare Committee of Université catholique de Louvain (Permit Number: 2013/UCL/MD/11 and 2017/UCL/MD/008). The day of vaginal plug was considered to be embryonic day (e) 0.5. A minimum of three embryos of the same genotype was analyzed in each experiment. The embryos were collected at e12.5 and e14.5. The *Hnf6;Oc2* and the *Pou2f2* mutant mice were previously described (Clotman et al., 2005, Corcoran et al., 1993, Jacquemin et al., 2000). In the *Hnf6^-/-^Oc2^-/-^* embryos, expression of *Oc3* is completely downregulated in the developing spinal cord (Kabayiza et al., 2017, Roy et al., 2012), enabling to study spinal cord development in the absence of the 3 OC factors. The mice and the embryos were genotyped by PCR (primer information available on request).

### *In situ* hybridization (ISH) and immunofluorescence labelings

For ISH, the collected embryos were fixed in ice-cold 4% paraformaldehyde (PFA) in phosphate buffered-saline (PBS) overnight at 4°C, washed thrice in PBS for 10 minutes, incubated in PBS/30% sucrose solution overnight at 4°C, embedded and frozen in PBS/15% sucrose/7.5% gelatin. Fourteen μm section were prepared and ISH was performed as previously described (Beguin et al., 2013, Francius et al., 2016, Pelosi et al., 2014) with DIG-conjugated Pou2f2 (NM_011138.1, nucleotides 604-1187) or Pou2f2 exon 5b (XM_006539651.3, nucleotides 643-876) antisense RNA probes. Control and Onecut mutant sections were placed adjacent on the same histology slides to minimize inter-slide variations in ISH signals.

For immunofluorescence, collected embryos were fixed in 4% PFA/PBS for 25 or 35 minutes according to their embryonic stage and processed as for ISH. Immunolabeling was performed on 14 μm serial cryosections as previously described (Francius and Clotman, 2010). Primary antibodies against the following proteins were used: Chx10 (sheep; 1:500; Exalpha Biologicals #X1179P), Foxp1 (goat; 1:1000; R&D Systems #AF4534), Gata3 (rat; 1:50; Absea Biotechnology #111214D02), GFP (chick; 1:1000; Aves Lab #GFP-1020), HNF6 (guinea pig; 1:2000; (Espana and Clotman, 2012b); or rabbit; 1:100; Santa Cruz #sc-13050; or sheep; 1:1000 R&D Systems #AF6277), cMaf (rabbit; 1:3000; kindly provided by H. Wende), MafA (guinea pig; 1:500; kindly provide by T. Müller), OC2 (rat; 1:400; (Clotman et al., 2005); or sheep; 1:500; R&D Systems #AF6294), OC3 (guinea pig; 1:6000; (Pierreux et al., 2004)), Pou2f2 (rabbit; 1:2000; Abcam #ab178679), Shox2 (mouse; 1:500; Abcam #ab55740), Sox1 (goat; 1:500; Santa Cruz #sc-17318). Secondary antibodies donkey anti-guinea pig/AlexaFluor 488, 594 or 647, anti-mouse/AlexaFluor 488, 594 or 647, anti-rabbit/AlexaFluor 594 or 647, anti-goat/AlexaFluor 488, anti-rat/AlexaFluor 647, anti-sheep/AlexaFluor 594 or 647, and goat anti-mouse IgG2A specific/AlexiaFluor 488, purchased from ThermoFisher Scientific or Jackson Laboratories were used at dilution 1:2000 or 1:1000, respectively.

### *In ovo* electroporation

*In ovo* electroporations were performed at stage HH14-16, as previously described (Roy et al., 2012). The coding sequence of the S_Pou2f2.4 transcript was amplified by overlapping-PCR using: forward 5’ GCTCTGTCTGCCCAAGAGAAA 3’ and reverse 5’ GTTGGGACAAGGTGAGCTGT 3’ primers for the 5’ sequence, forward 5’ CCACCATCACAGCCTACCAG 3’ and reverse 5’ ATTATCTCGAGCCAGCCTCCTTACCCTCTCT 3’ (designed to enable integration at the *Xho*I restriction site of the pCMV-MCS vector) primers for the 3’ sequence. This sequence was first subcloned in a pCR^®^II-Topo^®^ vector (Life Technologies, 45-0640) for sequencing then subcloned at the *Eco*RI (from the pCR^®^II-Topo^®^ vector) and *Xho*I restriction sites of a pCMV-MCS vector for the *in ovo* electroporation. The pCMV-Pou2f2 (0.5 μg/μl) vector was co-electroporated with a pCMV-eGFP plasmid (0.25μg/μl) to visualize electroporated cells. The embryos were collected 72 hours (HH27-28) after electroporation, fixed in PBS/4%PFA for 45 minutes and processed for immunofluorescence labelings as previously described (Francius and Clotman, 2010). To minimize stage and experimental condition variations, the non-electroporated side of the spinal cord was used as control for quantification and distribution analyses.

### Imaging and quantitative analyses

Immunofluorescence and ISH images of cryosections were acquired on an EVOS FL Auto Imaging System (Thermo Fisher Scientific) or a Confocal laser Scanning biological microscope

FV1000 Fluoview with the FV10-ASW 01.02 software (Olympus). The images were processed with Adobe Photoshop CS5 software to match brightness and contrast with the observation. Quantifications were performed by subtractive method (Francius and Clotman, 2010). For each embryo (*n* ≥ 3), one side of three to five sections at brachial, thoracic or lumbar level were quantified using the count analysis tool of Adobe Photoshop CS5 software. Raw data were exported from Adobe Photoshop CS5 software to Sigma Plotv12.3 software to perform statistical analyses. The histograms were drawn with Microsoft Excel. Adequate statistical tests (standard Student’s *t*-tests or Mann-Whitney U tests) were applied depending on the number of comparisons and on the variance in each group. Quantitative analyses were considered significant at *p* ≤ 0.05.

Quantitative analyzes of IN spatial distribution were performed as previously described (Kabayiza et al., 2017). Statistical analyses of ventral IN distribution were performed using a two-sample Hotelling’s T2, which is a two-dimensional generalization of the Student’s t test. The analysis was implemented using the NCSS software package.

### Microarray analyses

RNA was extracted from control or *Hnf6/Oc2* double-mutant spinal cords. The tissue was manually dissociated in Tripur isolation reagent (Roche, 11 667 165 001). After dissociation, chloroform (Merck Millipore, 1 02445 1000) was added to the sample, incubated at room temperature for 10 minutes and centrifugated for 10 minutes at 4°C. The aqueous phase was collected and the RNA was precipitated with isopropanol (VWR, 20880.310) and centrifugated for 15 minutes at 4°C. The pellet was washed in ethanol (Biosolve, 06250502) and centrifugated for 10 minutes at 4°C. The dried pellet was resuspended in RNAse free water. The integrity of the RNA was assessed using an Agilent RNA 6000 Nano assay procedure. For microarray analyzes, the RNA was converted in single-strand cDNA, labeled using the GeneChip^®^ WT PLUS Reagent Kit (Affymetrix) and hybridized on the GeneChip^®^ MoGene 2.0 ST array (Affymetrix, 90 2118) using Affymetrix devices: Genechip^®^ Fluidics Station 450, Genechip^®^ Hybridization oven 640, Affymetrix Genechip^®^ scanner and the Expression Consol software. The analyzes were performed using the R software. Microarray data have been deposited in the GEO repository (accession number: GSE117871).

### Amplification of *Pou2f2* isoforms and sequencing

Fragments of the different *Pou2f2* isoforms were amplified by RT-PCR from RNA of control B lymphocytes or embryonic spinal cords purified as described above using the iScript^™^ Reverse transcriptase and the 5x iScript^™^ reaction mix (BioRad). *Pou2f2* sequences (Table 2) were amplified using a GoTaq^®^ Green master mix (Promega, M712) or a Q5^®^ Hot Start High-Fidelity DNA Polymerase (New England BioLabs^®^ Inc, M0493S) (primer information available on request). Sequencing of the spinal *Pou2f2* exons was outsourced to Genewiz.

Quantitative real-time PCR was performed on 1/100 of the retrotranscription reaction using iTaq^™^ universal SYBR^®^ Green Supermix (BioRad, 172-5124) on a CFX Connect^™^ Real-Time System (BioRad) with the BioRad CFX Manage 3.1 software. Each reaction was performed in duplicate and relative mRNA quantities were normalized to the housekeeping gene RPL32 (primer information available on request). Relative expression changes between conditions were calculated using the ΔΔCt method. All changes are shown as fold changes.

## Acknowledgements

We thank members of the NEDI lab for material, technical support and discussions. We are grateful to Drs. T. Müller and H. Wende for kindly providing the guinea pig anti MafA and the rabbit anti cMaf antibodies, respectively, to Drs. F. Lemaigre and P. Jacquemin for the Hnf6/Oc2 double mutant mice, to Dr. C. Pierreux for the pCMV-MCS, to Dr. J. Ambroise for assistance with the microarray analyses and to Dr. L. Dumoutier for cDNA of B lymphocytes.

---

**Supplementary Figure S1.**
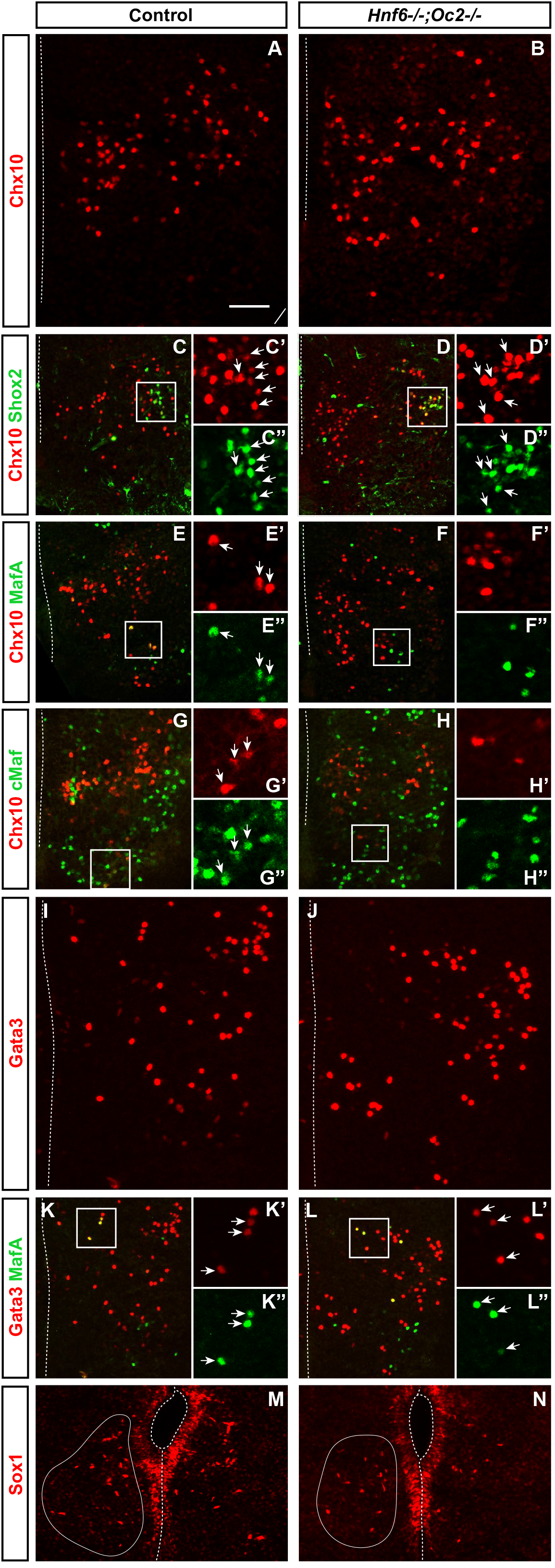
OC factors regulate the diversification of the V2 interneurons. **(A-H”)** Immunolabelings of the V2a generic marker Chx10 and markers of V2a subpopulations on transverse spinal cord sections (brachial or thoracic levels) control or *Hnf6^-/-^;Oc2^-/-^* mutant embryos at e14.5. Quantifications are shown in Fig 1. V2a **(A-B)**, Shox2+ V2a (arrows) and V2d interneurons (**C-D”**, arrowheads) are present in both control and double mutant embryos. **(E-E”)** MafA is present in a V2a supbopulation in control embryos (arrows), **(F-F”)** but is not detected in Chx10+ cells in the absence of OC factors. **(G-G”)** Similarly, cMaf is present in a subpopulation of V2a interneurons in control embryos (arrows), **(H-H”)** which is not the case in *Hnf6^-/-^;Oc2^-/-^* embryos. **(I-L”)** Immunolabelings of the V2b generic marker Gata3 and the marker of V2b subpopulation, MafA. V2b **(I-J”)** and MafA+ V2b (**K-L”**, arrows) interneurons are present in control and in *Hnf6^-/-^;Oc2^-/-^* embryos. **(M-N)** The number of V2c interneurons is unchanged is similarly detected in *Hnf6^-/-^;Oc2^-/-^* embryos as compared to control embryos (delineated cells). Sox1 in the ventricular zone labels neural progenitors. Scale bar = 50 μm.

**Supplementary Figure S2.**
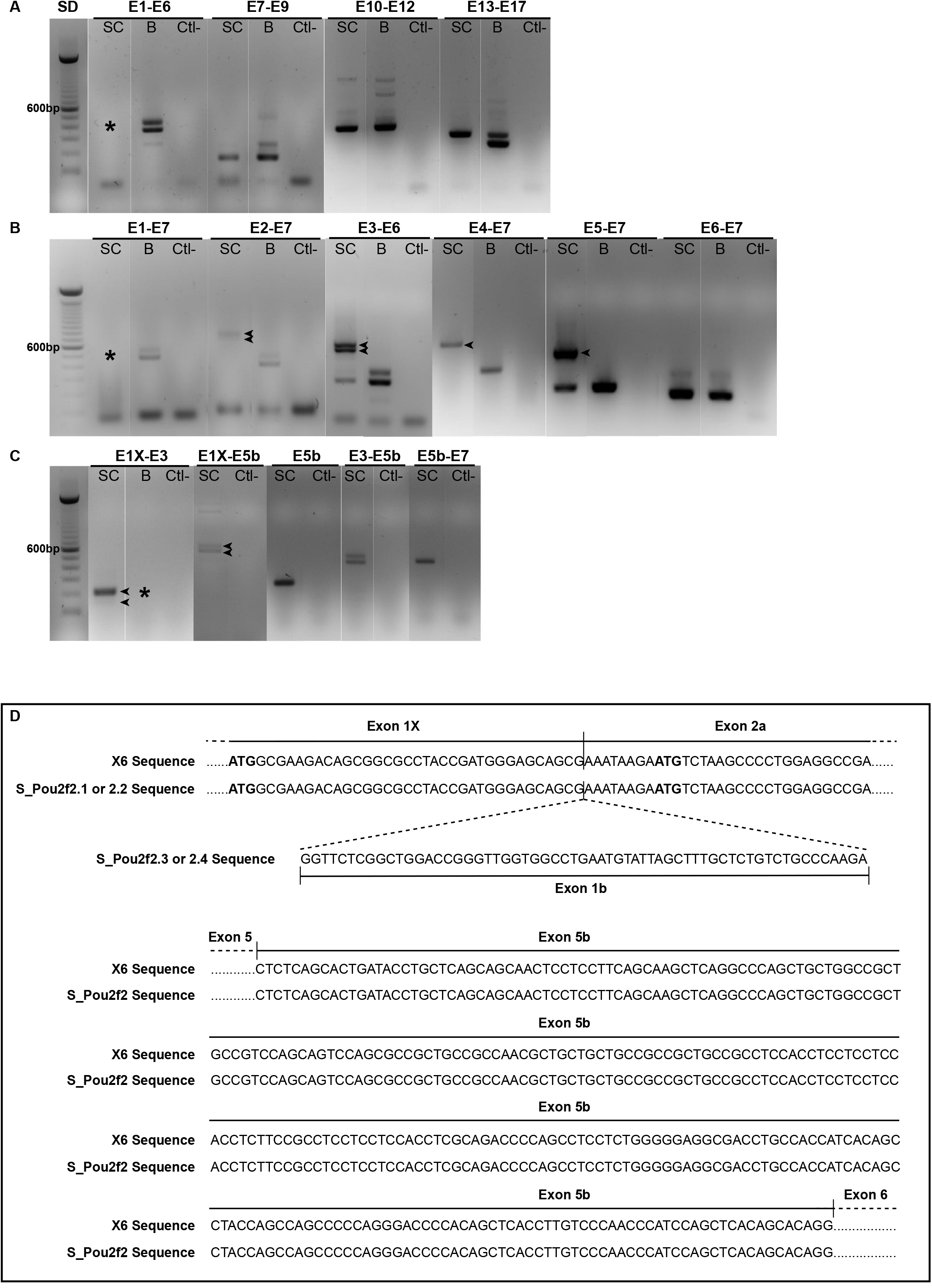
Pou2f2 RT-PCR experiments and sequencing of exon 1b and exon 5b of the spinal Pou2f2 isoforms. Composite assembly of electrophoresis images of RT-PCR amplification products for *Pou2f2* isoform sequences on embryonic spinal cord or B-cell RNA samples. Water was used as a negative control (Ctl-). **(A)** Amplifications from exon 1 (E1) to E6 on embryonic spinal cord RNA samples (asterisk) fail to amplify the RNA isoforms detected in B cell samples. In contrast, amplifications from E7 to E9, from E10 to E12 and from E13 to E17 show at least one similar amplicon in spinal cord samples and in B cell samples. **(B)** Amplification from E1 (other forward primer) to E7 also fails to amplify *Pou2f2* spinal cord isoforms. In contrast, amplifications from E2, 3, 4 and 5 to E7 produce systematically longer amplicons in spinal cord samples (arrowheads) as compared to B cell samples. The E6 to E7 amplification is similar in both samples. **(C)** Amplifications from E1X (present in the X6 sequence) to E3 do not produce amplicon in B-cell samples (asterisk). Amplifications from E1X to E3 or to E5b on spinal cord samples systematically produce 2 amplicons (arrowheads). Amplifications of the E5b exon sequence, from E3 to E5b and from E5b to E7 produce expected amplicons in spinal cord samples. **(D)** Comparison of exon 1b and exon 5b sequences in the predicted X6 sequence and the sequenced embryonic spinal cord *Pou2f2* isoforms. Sequences of E1X, E2a and E5b exons align (100% identity) with the predicted X6 sequence. Sequence of the additional alterative exon 1b is shown. SD = Size standard, E = Exon, SC = Embryonic spinal cord, B = B lymphocytes.

**Supplementary Figure S3.**
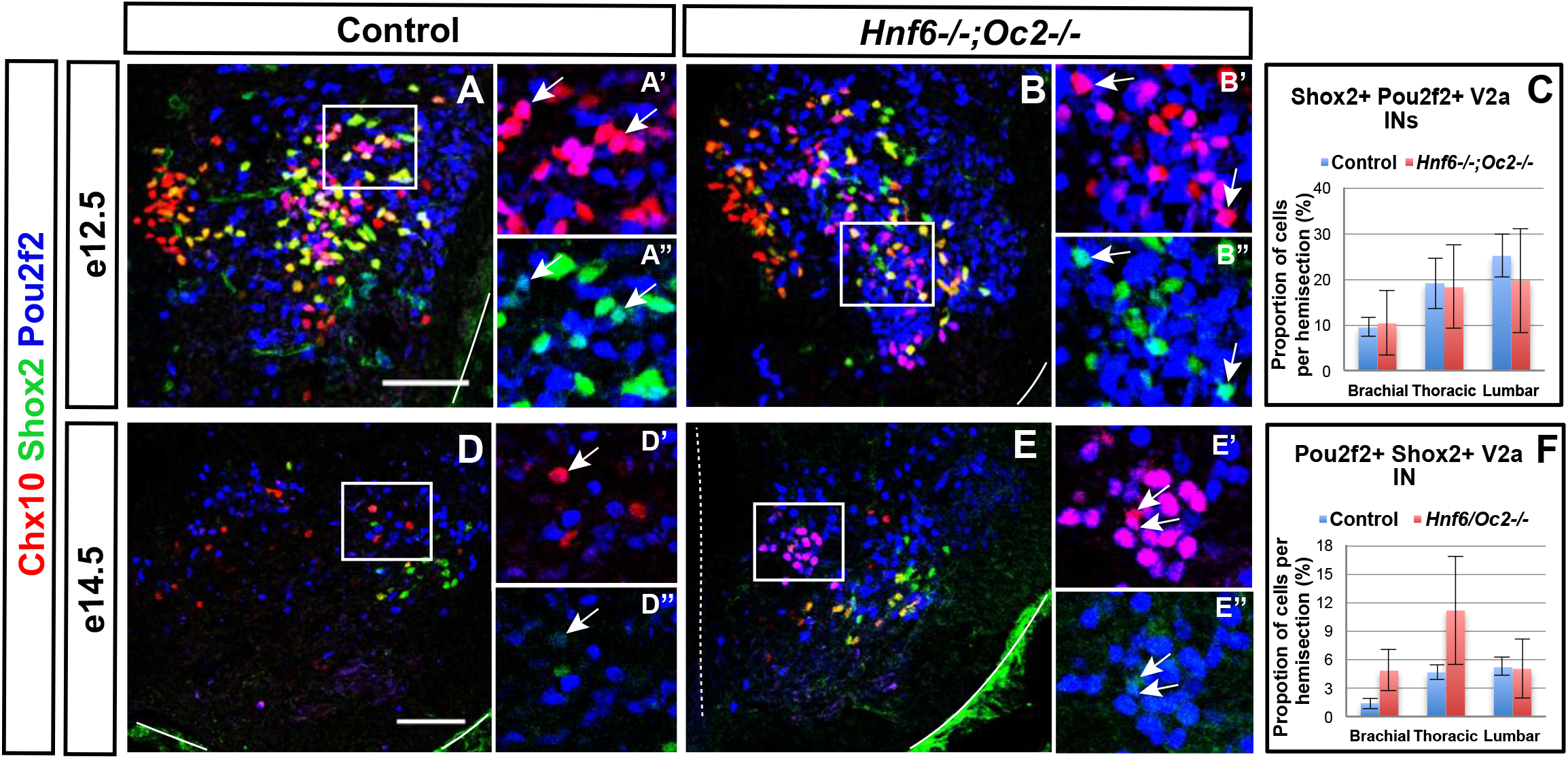
The number of Pou2f2+Shox2+ V2a interneurons is normal in *Hnf6^-/-^;Oc2^-/-^* mutant embryos. Immunolabelings on transverse spinal cord sections (brachial or thoracic levels) of control or *Hnf6^-/-^;Oc2^-/-^* mutant embryos. At e12.5 **(A-C)** and at e14.5 **(D-F)**, the number of V2a containing Shox2 and Pou2f2 is unchanged in the absence of OC factors. Mean values ± SEM. Scale bar = 50 μm.

**Supplementary Figure S4.**
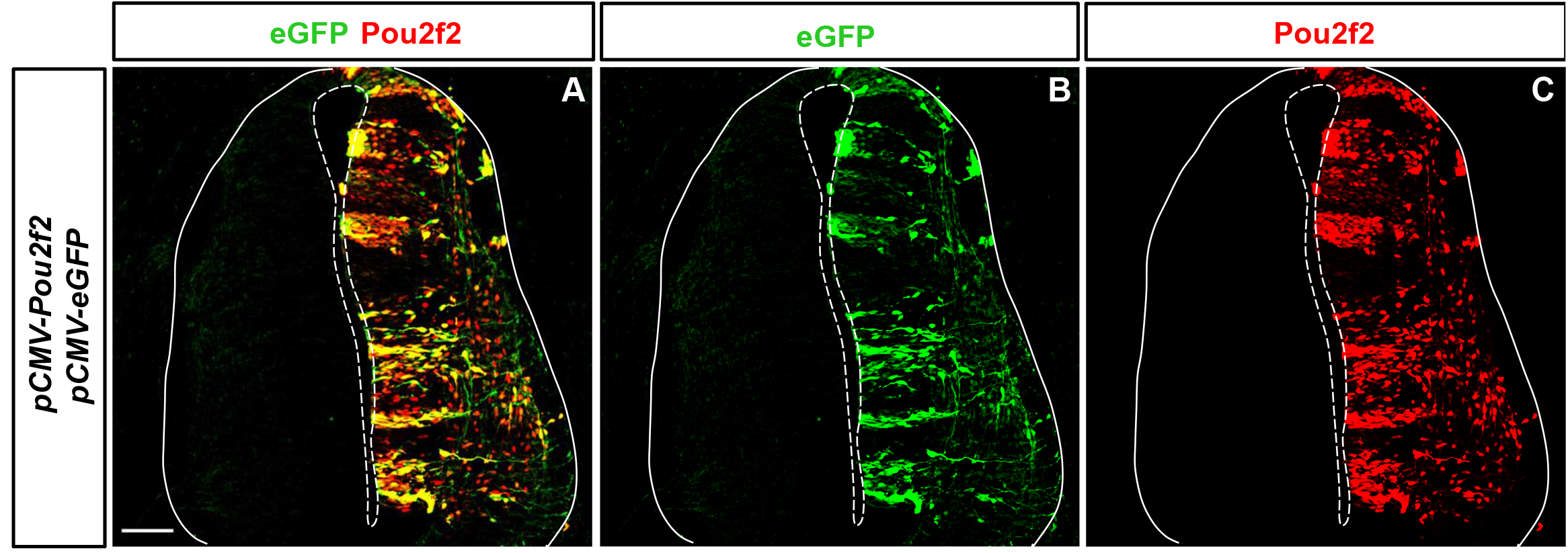
Efficacy of pCMV-eGFP and pCMV-Pou2f2 co-electroporation in the chicken embryonic spinal cord. **(A-C)** eGFP (green, **B**) and Pou2f2 (red, **C**) are present in a vast majority of cells along the dorso-ventral axis of the spinal cord. Scale bar = 50 μm.

**Supplementary Figure S5.**
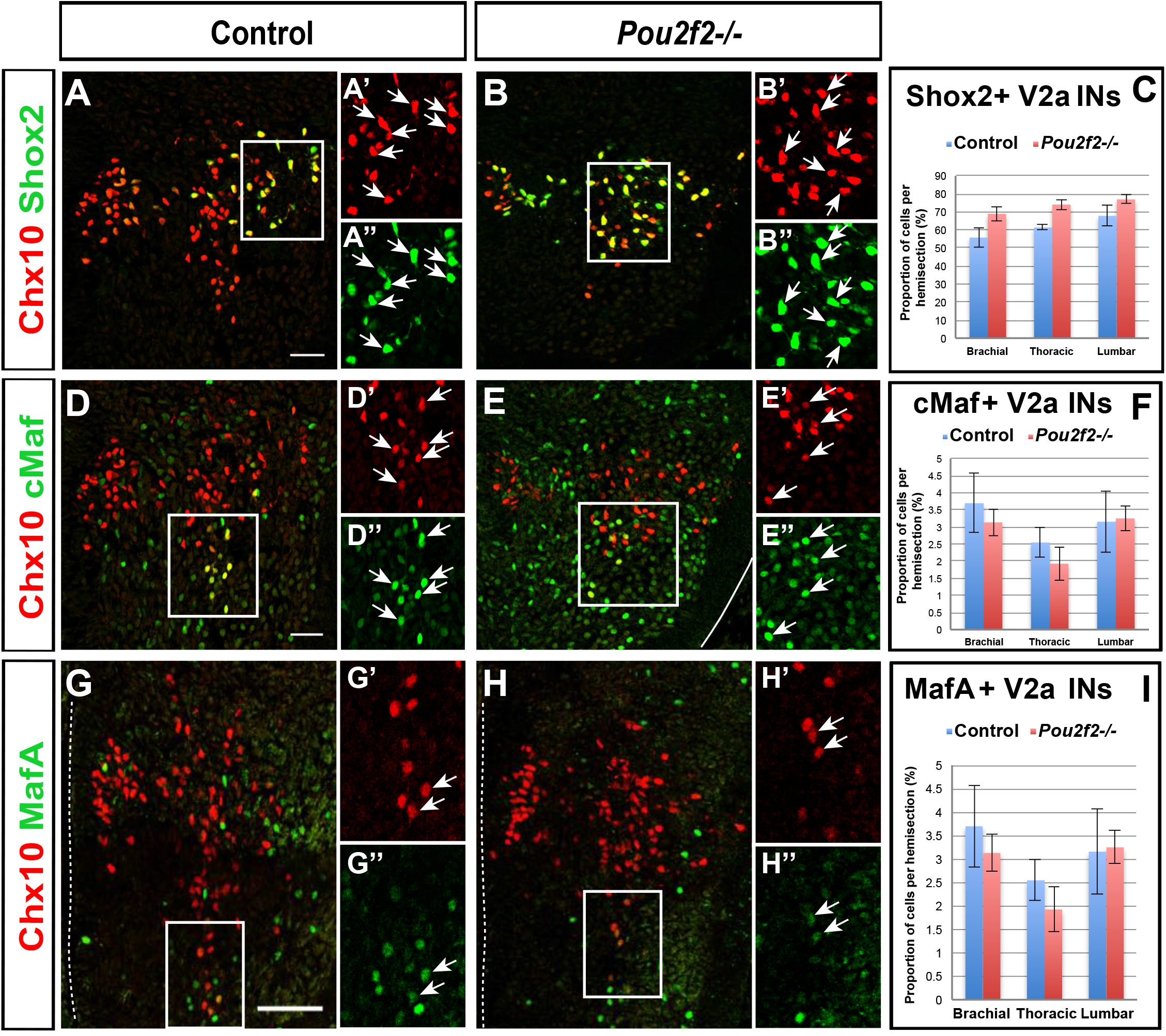
The number of MafA+ or cMaf+ V2a interneurons is normal in *Pou2f2^-/-^* mutant spinal cords. Immunolabelings on transverse spinal cord sections (brachial or thoracic levels) of control or *Pou2f2^-/-^* mutant embryos at e12.5. **(A-F)** Absence of Pou2f2 does not impact on the number of Shox2+ **(A-C)**, cMaf+ **(D-F)** or MafA+ **(G-I)** V2a interneurons. Mean values ± SEM. Scale bar = 50 μm.

**Supplementary Figure S6.**
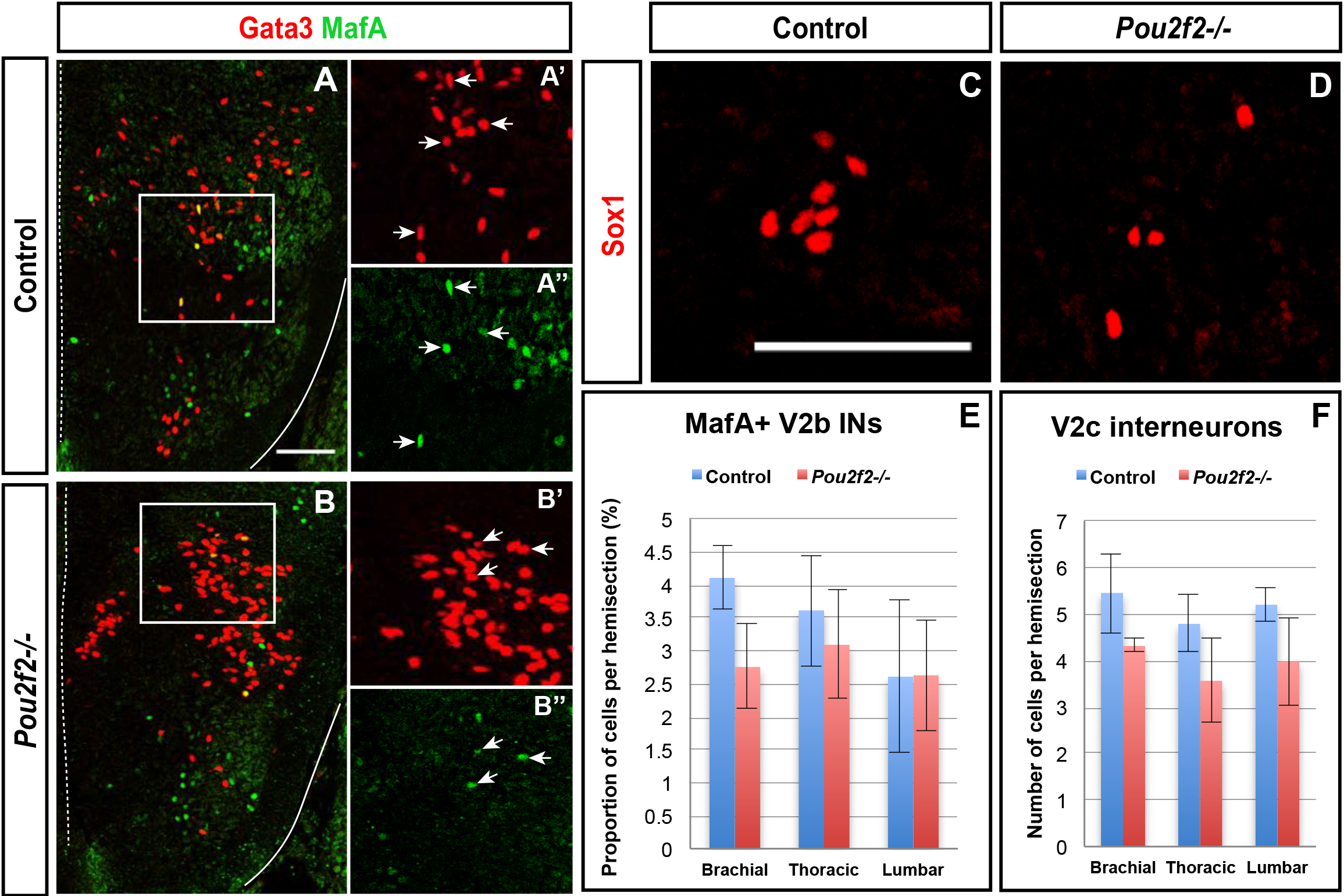
The number of MafA-positive V2b interneurons or of the V2c interneurons is normal in *Pou2f2^-/-^* mutant spinal cords. Immunolabelings on transverse spinal cord sections (brachial or thoracic levels) of control or *Pou2f2^-/-^* mutant embryos at e12.5. **(A-B, E)** The number of MafA+ V2b interneurons is not significantly altered in the absence of Pou2f2. **(C-D, F)** Similarly, the production of V2c interneurons is not affected in *Pou2f2* mutants. Mean values ± SEM. Scale bar = 50 μm.

